# *SURF1* mutations causative of Leigh syndrome impair human neurogenesis

**DOI:** 10.1101/551390

**Authors:** Gizem Inak, Agnieszka Rybak-Wolf, Pawel Lisowski, René Jüttner, Annika Zink, Barbara Mlody, Petar Glažar, Christopher Secker, Ummi H. Ciptasari, Werner Stenzel, Tobias Hahn, Sebastian Diecke, Josef Priller, Michael Gotthardt, Ralf Kühn, Erich E. Wanker, Nikolaus Rajewsky, Markus Schülke, Alessandro Prigione

## Abstract

Mutations in the mitochondrial complex IV assembly factor *SURF1* represent a major cause of Leigh syndrome (LS), a rare fatal neurological disorder. *SURF1*-deficient animals have failed to recapitulate the neuronal pathology of human LS, hindering our understanding of the disease mechanisms. We generated induced pluripotent stem cells from LS patients carrying homozygous *SURF1* mutations (SURF1 iPS) and performed biallelic correction via CRISPR/Cas9. In contrast to corrected cells, SURF1 iPS showed impaired neuronal differentiation. Aberrant bioenergetics in SURF1 iPS occurred already in neural progenitor cells (NPCs), disrupting their neurogenic potency. Cerebral organoids from SURF1 iPS were smaller and recapitulated the neurogenesis defects. Our data imply that *SURF1* mutations cause a failure in the development of maturing neurons. Using NPC function as an interventional target, we identified *SURF1* gene augmentation as a potential strategy for restoring neurogenesis in LS patients carrying *SURF1* mutations.

## Introduction

Leigh syndrome (LS, OMIM #256000) is a rare genetic neurological disorder characterized by symmetric lesions in the central nervous system (CNS), particularly in the basal ganglia and brain stem, and by psychomotor regression and lactic acidosis^1^. The symptoms typically manifest during the first year of life, with congenital onset occurring in one fourth of the cases^2^. The disease is fatal with a course that is variable but usually severe, and has a peak of mortality before the age of three^3^. LS is a genetically heterogeneous disease with pathogenic variants identified in more than 60 genes^4^. All of the affected genes are directly or indirectly involved in the maintenance of mitochondrial oxidative phosphorylation (OXPHOS), making LS one of the most prevalent mitochondrial disorders affecting 1/36,000 newborns^5^.

Recessive mutations in *SURF1* occur in about one third of all LS cases and represent the single most common cause of LS^6–8^. *SURF1* encodes the highly-conserved protein Surfeit locus protein 1 (SURF1), which is located in the inner mitochondrial membrane and is involved in the assembly of mitochondrial complex IV, i.e. the cytochrome C oxidase (COX)^9^. COX is the terminal enzyme of the electron transport chain catalyzing the transfer of electrons from cytochrome C to molecular oxygen. COX is composed of 13 subunits, three of which (MT-CO1, MT-CO2, MT-CO3) are encoded by the mitochondrial DNA (mtDNA) and constitute the hydrophobic catalytic core that is assembled first during formation of the complex^10^.

The molecular mechanisms underlying the neurological impairment caused by SURF1 defects in humans remain poorly understood. Consequently, there are currently no pathogenicity-based treatments available for affected patients^11,12^. An important obstacle in our understanding of the disease pathogenesis is the paucity of adequate model systems^13^. *SURF1* knock-out mice failed to recapitulate the neurological phenotypes seen in humans and even led to prolonged lifespan, despite the presence of COX defects and mild lactic acidosis^14,15^. In *Drosophila melanogaster*, CNS-specific knock-down of *SURF1* increased longevity and did not produce neurological defects, while constitutive *SURF1* knock-down resulted in embryonic lethality^16^. *SURF1* knock-down in zebrafish caused reduced COX activity and developmental defects in endodermal tissue with a neurological impairment mainly occurring at the level of motoneurons and the peripheral nervous system^17^. *SURF1* knock-out in piglets resulted in a severe lethal phenotype but the neurological defects were mild, mainly comprising a slight delay of CNS development without lactic acidosis and without apparent COX deficiency^18^.

In order to dissect the molecular mechanisms underlying the neuronal pathology in LS patients carrying homozygous *SURF1* mutations (LS^SURF1^), we generated induced pluripotent stem cells from LS^SURF1^ patients (SURF1 iPS). Neuronal cultures and cerebral organoids derived from SURF1 iPS exhibited features indicative of impaired neurogenesis. Bioenergetic dysfunction occurred already at the level of neural progenitor cells (NPCs), and led to defects in neuronal generation and maturation, disrupting the branching capacity and firing activity of neurons. Biallelic correction of *SURF1* mutation *via* CRISPR/Cas9 genome editing reverted the phenotypes. Our data imply that LS^SURF1^ causes a failure in the development of maturing neurons. We employed NPC function as an interventional target and discovered that wild-type SURF1 over-expression was effective in improving NPC bioenergetics and promoting neurogenesis. Altogether, we propose aberrant neurogenesis as a central pathogenetic mechanism in LS^SURF1^ and suggest a potential strategy for restoring neuronal function in this fatal pediatric disease.

## Results

### Genetic repair in SURF1 iPS restores COX activity

We obtained skin fibroblasts from two LS^SURF1^ patients belonging to two distinct consanguineous families: S1A carrying the homozygous mutation c.530T>G (p.V177G), and S1C carrying the homozygous mutation c.769G>A (p.G257R) (**Fig. S1**). We used Sendai virus-based reprogramming to generate the respective SURF1 iPS lines SVS1A and SVS1C, which we confirmed to be pluripotent (**Fig. 1a; Fig. S2a-b; Fig S2e**) while still harboring the parental *SURF1* mutations on both alleles (**Fig. 1a; Fig. S2c-d**).

**Fig. 1.**
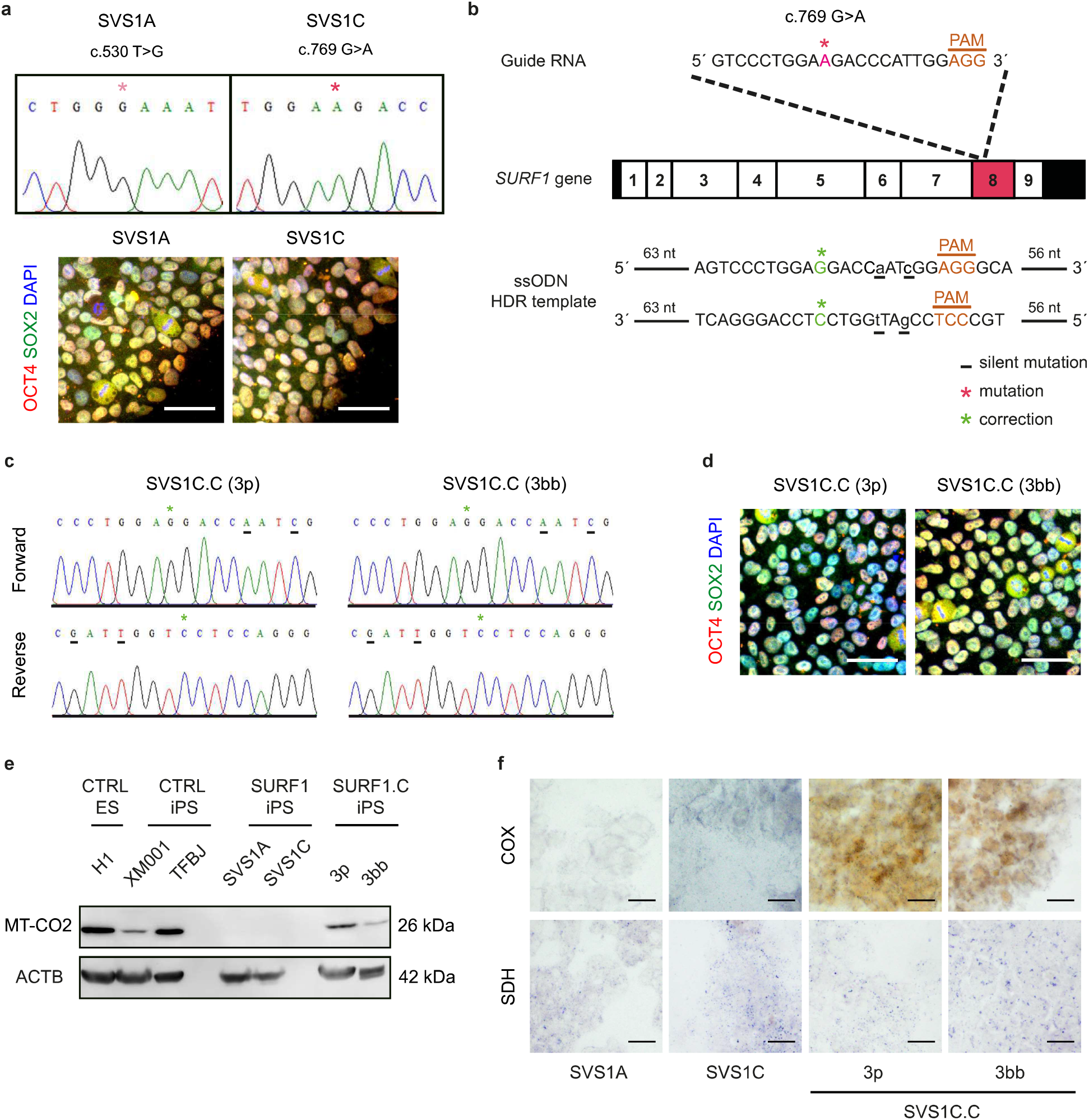
CRISPR/Cas9-based correction of iPSCs derived from LS^SURF1^ patients. **a,** Upper panel: Sanger sequencing confirmation of the LS^SURF1^-iPSCs SVS1A and SVS1C carrying the patient-specific homozygous mutations c.530T>G (p.Vl77G) and c.769G>A (p.G257R), respectively. Lower panel: iPSC lines expressed pluripotency-associated marker proteins OCT4 and SOX2. Scale bar: 50 μm. **b**, CRISPR/Cas9 strategy for repairing the *SURF1* c.769G>A mutation on both alleles in SVS1C. ssODNs: single-stranded oligodeoxynucleotides; HDR: homologous direct repair, **c**, Sanger sequencing confirmation of genetically repaired iPSCs SVS1C.C (clones 3p and 3bb). **d**, Corrected iPSC lines retained pluripotent identity. Scale bar: 50μm. **e**, Representative immunoblot analysis for MT-CO2 in iPSC-derived NPCs from CTRL ES (HI), CTRL iPS (XM001 and TFBJ), SURF1 iPS (SVS1A and SVS1C), and genetically corrected SVS1C.C (clones 3p and 3bb). Protein levels were normalized to β-actin (ACTB). Similar results were obtained in n = 2 independent experiments, **f**, Mitochondrial enzyme activity in NPCs derived from SVS1A, SVS1C, and SVS1C.C (clones 3p and 3bb). Activity of complex IV cytochrome oxidase (COX) was detectable by a positive color reaction (brown) only in corrected SVS1C cells, while the activity of complex II succinate dehydrogenase (SDH, blue coloration) was present in all cells. Similar results were obtained in n = 2 independent experiments.

We corrected the c.769G>A mutation on both alleles in SVS1C using clustered regularly interspaced short palindromic repeats (CRISPR)/Cas9 and obtained SVS1C-corrected cells (SVS1C.C) (**Fig. 1b-d; Fig. S3a-b**). We achieved genetic correction using high-fidelity eCas9, which has limited off-target effects^19^, and by including plasmids expressing RAD52 and dominant negative p53-binding protein 1 (dn53BP1) in order to promote homologous direct repair (HDR) and inhibit non-homologous end joining (NHEJ)^20^ (**Fig. S3a**). In the HDR template, we introduced two silent mutations that we could trace in the sequences of the SVS1C.C clones (3p and 3bb) to confirm the faithful correction of the mutation (**Fig. 1b-c**). The corrected lines retained features of pluripotency (**Fig. 3d; Fig. S3c**) and had normal karyotypes (**Fig. S3e**).

We monitored COX activity in NPCs derived from SURF1 iPS (SVS1A and SVS1C) and from genetically corrected iPSCs (SVS1C.C, clones 3p and 3bb). We determined the protein abundance of the MT-CO2 subunit, a key component of the COX enzyme catalytic core, and the enzymatic activity of the entire COX complex. In NPCs from SVS1A and SVS1C, MT-CO2 protein was almost undetectable (**Fig. 1e**) and COX activity was very low (**Fig. 1f**). In contrast, NPCs derived from CRISPR/Cas9-corrected SVS1C.C lines reexpressed MT-CO2 (**Fig. 1e**) and had their COX activity restored (**Fig. 1f**). The activity of complex II succinate dehydrogenase (SDH) remained similar in all NPCs (**Fig. 1f**), confirming that the defects in *SURF1*-mutant cells were specific for complex IV. Overall, the findings demonstrated that the mutation repair in SVS1C.C had led to functional recovery of SURF1-mediated COX assembly.

### *SURF1* mutations impair the generation of neuronal cells

In order to address the influence of *SURF1* mutations on the development and function of human neurons, we differentiated all induced pluripotent stem cells (iPSCs) into functional neuronal cultures that were enriched for midbrain dopaminergic (DA) neurons^21^ (**Figs. 2a; S4a-c**), as degeneration of DA neurons is thought to cause the predominant basal ganglia pathology seen in LS patients (**Fig. S1a-f**)^22^ Besides neurons, the cultures contained mature astrocytes (**Fig. 2a**), which are important CNS sources of lactate^23^, whose levels are increased in the liquor and serum of most LS patients^11^.

**Fig. 2.**
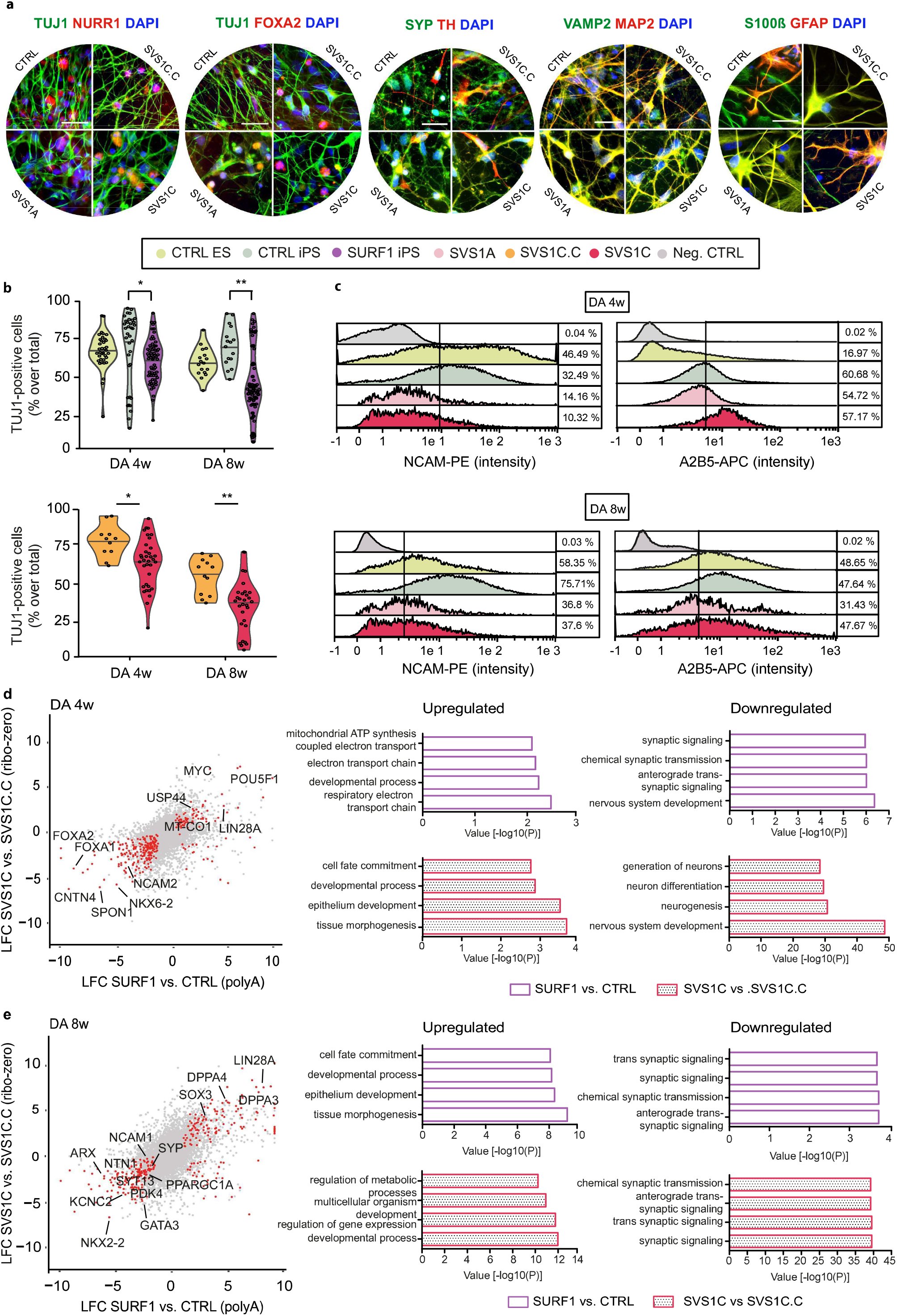
Impaired neurogenesis in *SURF1*-mutant iPSCs. **a,** Dopaminergic (DA)-enriched neuronal cultures from CTRL iPS (TFBJ and XM001), SURF1 iPS (SYS1A and SVS1C), and genetically corrected SVS1C.C expressed the neuronal marker TUJ1, the synaptic markers SYP and VAMP2, the DA markers NURR1, FOXA2, and TH, and the astrocyte markers GFAP and S100ß. Scale bar: 50 μm. **b**, Upper panel: high-content analysis (HCA)-based quantification of TUJ1-expressing neurons in 4w DA cultures (n = 4 independent experiments) and 8w DA cultures (n = 2 independent experiments) derived from CTRL ES (HI), CTRL iPS (TFBJ) and SURF1 iPS (SVS1A and SVS1C). Violin plots show median and the kernel probability density (* p < 0.05, ** p < 0.01; one-way ANOVA followed by Tukey Honestly Significant Differences). Lower panel: TUJ1 quantification in 4w and 8w DA cultures from SVS1C.C and SVS1C. Violin plots show median and the kernel probability density (* p < 0.05, ** p < 0.01; Mann-Whitney *U* test), **c**, Representative Magnetic-Activated Cell sorting (MACs)-based histogram plots of NCAM-positive cell population and A2B5-positive cell population in 4w DA cultures (upper panel) and 8w DA culture (lower panel) from CTRL ES (HI), CTRL iPS (XM001) and SURF1 iPS (SVS1A and SVS1C). Fluorescence intensity (x-axis) was plotted against relative cell number (y-axis). Percentage values refer to the number of positive cells over the total number of cells. Similar results were obtained in n = 2 independent experiments, **d-e**, Transcriptomics of 4w and 8w DA cultures. Left: Plots showing log fold changes (LFC) of genes in SVS1C *vs.* SVS1C.C in relation to LFC of genes in SURF1 iPS (SVS1A and SVS1C) *vs.* CTRL (HI and XM001). Highlighted genes were among the top consensually downregulated and upregulated genes across the two datasets. Right: top-four most significantly biological processes (gene ontology terms) that were upregulated or downregulated in SURF1 iPS (SVS1A and SVS1C) in comparison to CTRL (HI and XM001) (empty bars) or in SVS1C in comparison to SVS1C.C (dotted bars).

We assessed DA cultures at 4 weeks (4w) and 8 weeks (8w) after differentiation from NPCs. We found a reduction in the numbers of neurons in DA cultures obtained from SURF1 iPS as compared to DA cultures derived from healthy induced pluripotent stem cells (CTRL iPS) or from human embryonic stem cells (CTRL ES) (**Fig. 2b**). Mutation correction in SVS1C.C restored neuronal differentiation (**Fig. 2b**), confirming that *SURF1* mutations specifically impaired neurogenesis. DA cultures from SVS1A and SVS1C showed a reduced amount of NCAM-positive neuron-restricted progenitors compared to cultures from CTRL ES and CTRL iPS (**Fig. 2c**). Conversely, the number of A2B5-positive glial-restricted progenitors did not change (**Fig. 2c**). The number of cells positive for the astrocyte marker GLAST was also unchanged (**Fig. S4d-f**). Astrocyte-derived cytokines^24^ increased over time in mature cultures but did not exhibit significant differences between LS^SURF1^ patients and controls (Table S1). Taken together, *SURF1* mutations affected neuronal production possibly through defects of neurogenesis arising at the level of neuron-committed NPCs.

We performed transcriptome analysis of 4w and 8w DA cultures from SURF1 iPS, CTRL iPS, and CTRL ES using polyA mRNA-sequencing (**Fig. S5a**) and of 4w and 8w DA cultures from SVS1C and SVS1C.C using ribo-zero total RNA-seq (**Fig. S5a**). There was a strong correlation between the two datasets and we focused on the genes and pathways that were consensually differentially regulated in both settings. Downregulated genes in *SURF1-* mutant 4w DA cultures included genes involved in neurogenesis and nervous system development such as *NKX2-2, FOXA1, FOXA2*, and *NCAM2* (**Fig. 2d; Fig S5b; Table S2; Table S4**), as well as genes important for axonal outgrowth and synaptic signaling like *SPON1* and *CNTN4* (**Fig. 2d; Fig. S5c-d; Table S2; Table S4**). The downregulation of genes regulating synaptic function *(SYP, SYT13, KCNC2)* and axon guidance *(ARX, NTN1)* in *SURF1*-mutant cultures became even more evident at 8 weeks of differentiation (**Fig. 2e; Fig. S5b-d; Table S3; Table S4**). At the same time, *SURF1*-mutant DA cultures at 4w and 8w upregulated genes associated with embryonic development and pluripotency such as *MYC, LIN28A, POU5F1, USP44, DPPA3*, and *DPPA4* (**Fig. 2d-e; Fig. S5e; Table S2-S4**), and genes associated with neural progenitor cell fate identity like SOX3 (**Fig. 2e**). Overall, *SURF1* mutations caused a transcriptional reconfiguration suggestive of a suppression of neuronal differentiation and maturation which was accompanied by a failure to repress the pluripotent and precursor fate identities.

### Impairment of bioenergetics in *SURF1*-mutant cells already at the level of neural progenitors

The transcriptional signature of SURF1-mutant 4w DA cultures indicated an upregulation of mtDNA-encoded genes (such as *MT-CO1)* (**Fig. 2d; Fig. S6a; Table S2; Table S4**). At the beginning of differentiation, neuronal cells have increased energy demands and may thus attempt to compensate their COX deficiency through augmented expression of components of the catalytic core of COX enzyme. However, upon prolonged differentiation this compensatory mechanism may be shut down as it becomes insufficient. Accordingly, 8w neuronal cultures from SURF1 iPS downregulated *PPRGC1A (PGC1A)* (**Fig. 2e; Fig. 3a; Fig. S6a; Table S3; Table S4**), a master activator of mitochondrial biogenesis and OXPHOS metabolism that is important for neuronal maturation^25,26^. SURF1 iPS 8w DA cultures also downregulated *PDK3* and *PDK4* (**Fig. 2e; Fig. 3a; Fig. S6a; Table S3; Table S4**), which are negative regulators of glucose oxidation and crucial for maintaining energy homeostasis^27,28^. At the same time, 4w and 8w DA cultures from SURF1 iPS showed increased expression of glycolysis-promoting lactate dehydrogenase enzymes *LDHA* and *LDHD* (**Fig. 3a; Fig. S6a**). Taken together, maturing neuronal cultures carrying *SURF1* mutations might fail to activate mitochondrial metabolism over time and therefore retain features of glycolytic metabolism.

**Fig. 3.**
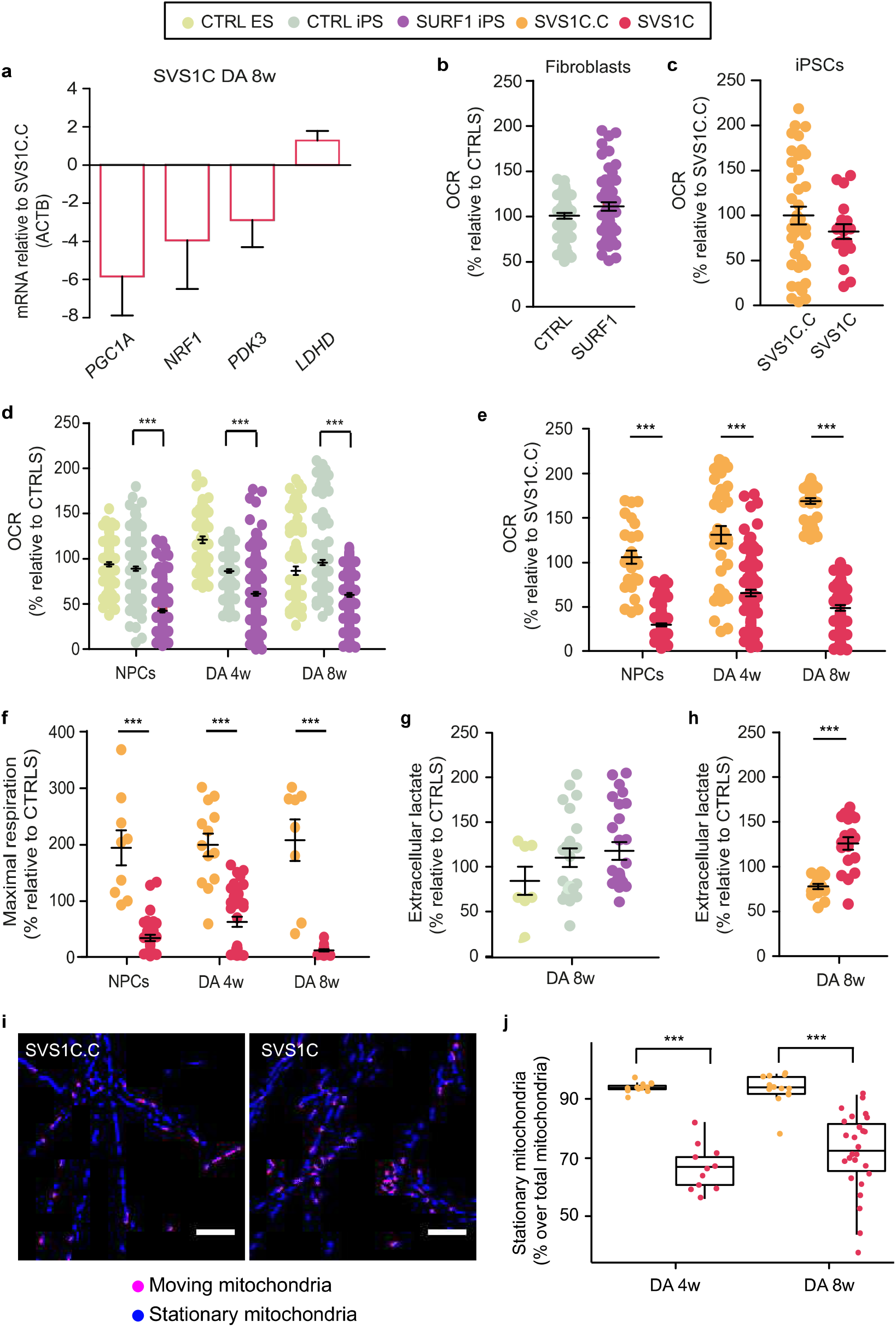
SURF1 defects impair mitochondrial bioenergetics in NPCs and differentiated neurons. **a,** Quantitative real-time RT-PCR showing the downregulation of genes promoting mitochondrial oxidative bioenergetics and the upregulation of a glycolytic-promoting gene in 8w DA cultures from SVS1C in comparison to SVS1C.C. Data were normalized to *ACTB* (mean +/− s.d; n = 3 independent experiments), **b**, Oxygen consumption rate (OCR) assessed via Seahorse extracellular flux analysis in fibroblasts from healthy CTRL (CON1 and BJ) and SURF1 patients (S1A and SIC) (mean +/− s.e.m.; n = 2 independent experiments; Mann-Whitney *U* test, not significant), **c**, OCR in iPSC lines SVS1C.C and SVS1C (mean +/− s.e.m.; Mann-Whitney *U* test; not significant), **d**, OCR in NPCs (n = 5 independent experiments), 4w DA cultures (n = 3 independent experiments), and 8w DA cultures (n = 3 independent experiments) from CTRL ES (HI), CTRL iPS (XM001), and SURF1 iPS (SVS1A and SVS1C) (mean +/− s.e.m.; *** p < 0.001; one-way ANOVA followed by Bonferroni multiple comparison test), **e-f**, OCR and maximal respiration rate in NPCs and 4w and 8w DA cultures from SVS1C.C and SVS1C (mean +/− s.e.m.; *** p < 0.001, Mann-Whitney *U* test), **g**, Quantification of extracellular lactate in 8w DA cultures from CTRL ES (HI), CTRL iPS (XM001), and SURF1 iPS (SVS1A and SVS1C) (mean +/− s.e.m.; n = 4 independent experiments; one-way ANOVA followed by Bonferroni multiple comparison test; not significant), **h**, Quantification of extracellular lactate in 8w DA cultures from SVS1C.C and SVS1C (mean +/− s.e.m.; n = 3 independent experiments; *** p < 0.001; Mann-Whitney *U* test), **i**, Representative images of 8w DA cultures live-stained with MitoTrackerRed in SVS1C.C (total mitochondria = 87, moving mitochondria = 45) and SVS1C (total mitochondria = 76, moving mitochondria = 26). See also Supp. videos 1-4. Scale bar: 10 μm. **j**, Stationary mitochondria in 4w and 8w DA cultures from SVS1C.C and SVS1C (horizontal bars indicate the median, boxes are defined by the first and third quartiles, whiskers span 1.5 x inter-quartile range; *** p < 0.001, Mann-Whitney *U* test).

In order to dissect the functional consequences of *SURF1* mutations on cellular bioenergetics, we investigated the oxygen consumption rate (OCR) in living cells. We did not observe differences between LS^SURF1^ patients and controls at the level of somatic fibroblasts (**Fig. 3; Fig. S6b; Fig S6i**) or iPSCs (**Fig. 3c; Fig. S6c; Fig. S6j**). On the other hand, *SURF1-* mutant neural cells exhibited significantly reduced mitochondrial respiration (basal OCR, maximal respiration, and ATP production) (**Fig. 3d; Fig. S6d-g**). Mitochondrial bioenergetics was restored upon mutation correction in SVS1C.C-derived neural cells (**Fig. 3e-f; Fig. S6d-f; Fig. S6h**). Bioenergetic defects were present throughout neuronal differentiation but became biochemically apparent already at the level of NPCs (**Fig. 3d-f; Fig. S6d; Fig. S6g-h**). When compared to isogenic control cultures, 8w DA cultures from SVS1C produced an increased amount of glycolysis-derived lactate, a pathognomonic feature of LS (**Fig. 3g-h**). Overall, the bioenergetic assessments supported our observation that *SURF1* mutations interfere with the function of NPCs, which may in turn affect the functionality of differentiated neurons.

We assessed the consequences of the bioenergetic defects in differentiated neurons through quantification of mitochondrial movement. SVS1C-derived DA cultures showed an abnormal increase in mitochondrial motility in comparison to SVS1C.C cultures (**Fig. 3i; Suppl. video 1-4**), causing a significantly reduced number of stationary mitochondria (**Fig. 3j**). Stationary mitochondria are crucial for axonal branching and synaptic homeostasis^29,30^. Hence, the disruption of mitochondrial bioenergetics and motility in SURF1-mutant cultures might negatively affect the branching and firing capacity of differentiating neurons.

### *SURF1* mutations disrupt neuronal branching and activity

We investigated whether the defects observed in *SURF1*-mutant NPCs altered not only the number of neurons and their metabolic properties, but also in their firing capacity. Patch-clamp analysis demonstrated that differentiating neurons from controls and patients matured over time, as indicated by their passive membrane properties showing time-dependent increase in cell capacitance (as neurons increase in size) (**Fig. S7a**) and decrease in input resistance (as neurons acquire more ion channels in their plasma membrane) (**Fig. S7b**). However, the active membrane properties of neurons (i.e. electrical potentials across the plasma membrane activated by voltage, ligands, or second messengers) differed between patients and controls. We observed a significant increase in sodium and potassium currents at 8w compared to 4w in neurons derived from CTRL iPS (**Fig. 4a**) and from CTRL ES (**Fig S7d**). This increase in currents at 8w was less pronounced in neurons differentiated from SURF1 iPS (**Fig. 4b**). Neurons from SURF1 iPS also contained a higher number of non-spikers and a lower number of multiple-spikers as compared to neurons from CTRL iPS (**Fig. 4b-c; Fig. S7c; Fig. S7e-f**). Genetic correction reconstituted full neuronal activity, as SVS1C.C-derived DA cultures exhibited significantly increased currents as compared to SVS1C-derived cultures at 4w (**Fig. 4d**) and at 8w (**Fig. 4e**). Neurons from SVS1C.C also contained a lower number of non-spikers and a higher number of multiple-spikers than neurons from SVS1C at 4w (**Fig. S7c**) and at 8w (**Fig. 4f; Fig S7e**). Moreover, SVS1C.C neurons exhibited post-synaptic currents indicative of synaptic activity, which we could not observe in SVS1C neurons (**Fig. 4g**).

**Fig. 4.**
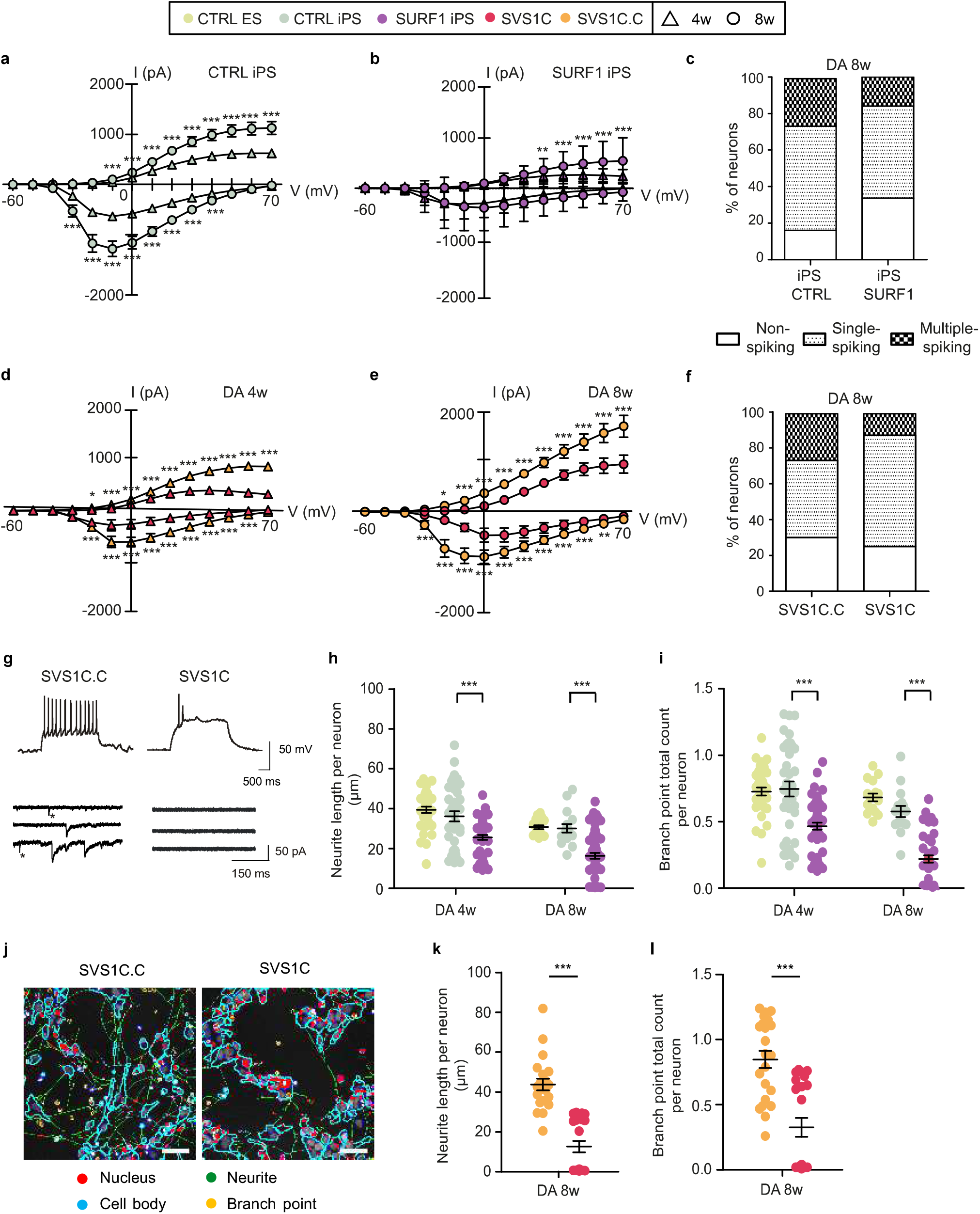
Aberrant neuronal firing and branching in LS^SURF1^ neurons. **a,** Patch-clamp measurement of sodium and potassium currents in 4w and 8w DA cultures from CTRL iPS (XM001 and TFBJ) (mean +/− s.d.; n = 7 independent experiments; *** p < 0.001; two-way ANOVA). **b**, Sodium and potassium currents in 4w and 8w DA cultures from SURF1 iPS (SVS1A and SVS1C) (mean +/− s.d.; n = 4 independent experiments; ** p < 0.01, *** p < 0.001; two-way ANOVA). **c**, Percentage of neurons (mean) that are not-spiking, single-spiking, or multiple spiking in 8w DA cultures from CTRL iPS (XM001 and TFBJ) and SURF1 iPS (SVS1A and SVS1C) (n = 4 independent experiments), **d**, Sodium and potassium currents in 4w DA cultures from SVS1C.C and SVS1C (mean +/− s.d.; n = 2 independent experiments; * p < 0.05, *** p< 0.001; two-way ANOVA). **e**, Sodium and potassium currents in 8w DA cultures from SVS1C.C and SVS1C (mean +/− s.d.; n = 2 independent experiments; * p < 0.05, ** p < 0.01, *** p < 0.001; two-way ANOVA). **f**, Percentage of neurons (mean) that are not-spiking, single-spiking, or multiple spiking in 8w DA cultures from SVS1C.C and SVS1C (n = 2 independent experiments), **g**, Representative electrophysiology traces showing that 8 w DA cultures from SVS1C.C contained neurons showing multiple spiking activity and post-synaptic currents, **h-i**, HCA-based quantification of neuronal branching complexity in 4w and 8w DA cultures from CTRL ES (HI), CTRL iPS (XM001), and SURF1 iPS (SVS1A and SVS1C) (mean +/− s.e.m.; n = 3 independent experiments; *** p< 0.001; one-way ANOVA followed by Bonferroni multiple comparison test), **j**, Representative images of the HCA mask for neuronal profiling showing decreased number of branches (green) and branching points (yellow) in 8w DA cultures from SVS1C.C compared to SVS1C. Scale bar: 50 μm. **k-1**, Neuronal profiling in 8w DA cultures from SVS1C.C and SVS1C (mean +/− s.e.m.; n = 3 independent experiments; *** p< 0.001; Mann-Whitney Ł/test).

We next addressed the capacity of neurons to generate branches using high-content analysis (HCA)-based quantification of neurite length per neuron and branching points per neuron. Neurite outgrowth was significantly lower in neurons from SURF1 iPS than in neurons from CTRL iPS (**Fig. 4h-i**). This was particularly evident in 8w DA cultures. Correction of *SURF1* mutation in SVS1C.C neurons restored the neuronal complexity in neurites and branching patterns (**Fig. 4j-l**). Altogether, *SURF1* mutations impaired the functionality of maturing neurons by disrupting their firing properties and their ability to establish synaptic connections.

### Neurogenesis defects in *SURF1*-mutant cerebral organoids

Since monolayer cultures do not allow studying the manifestation of spatial defects, we employed a three-dimensional (3D) model of neuronal development (**Fig. S8a-b**) to explore the consequences of *SURF1* mutations on organogenesis. Cerebral organoids derived from SVS1C were significantly smaller than organoids differentiated from the isogenic control SVS1C.C (clones 3p and 3bb) (**Fig. 5a-b; Fig. S8c**). SVS1C organoids exhibited a dramatic disruption of the neuroepithelial layer surrounding fluid-filled cavities that are reminiscent of embryonic ventricles (**Fig. 5c; Fig. S8d**). These macroscopic phenotypes pointed towards neurodevelopmental defects caused by impaired neurogenesis.

**Fig. 5.**
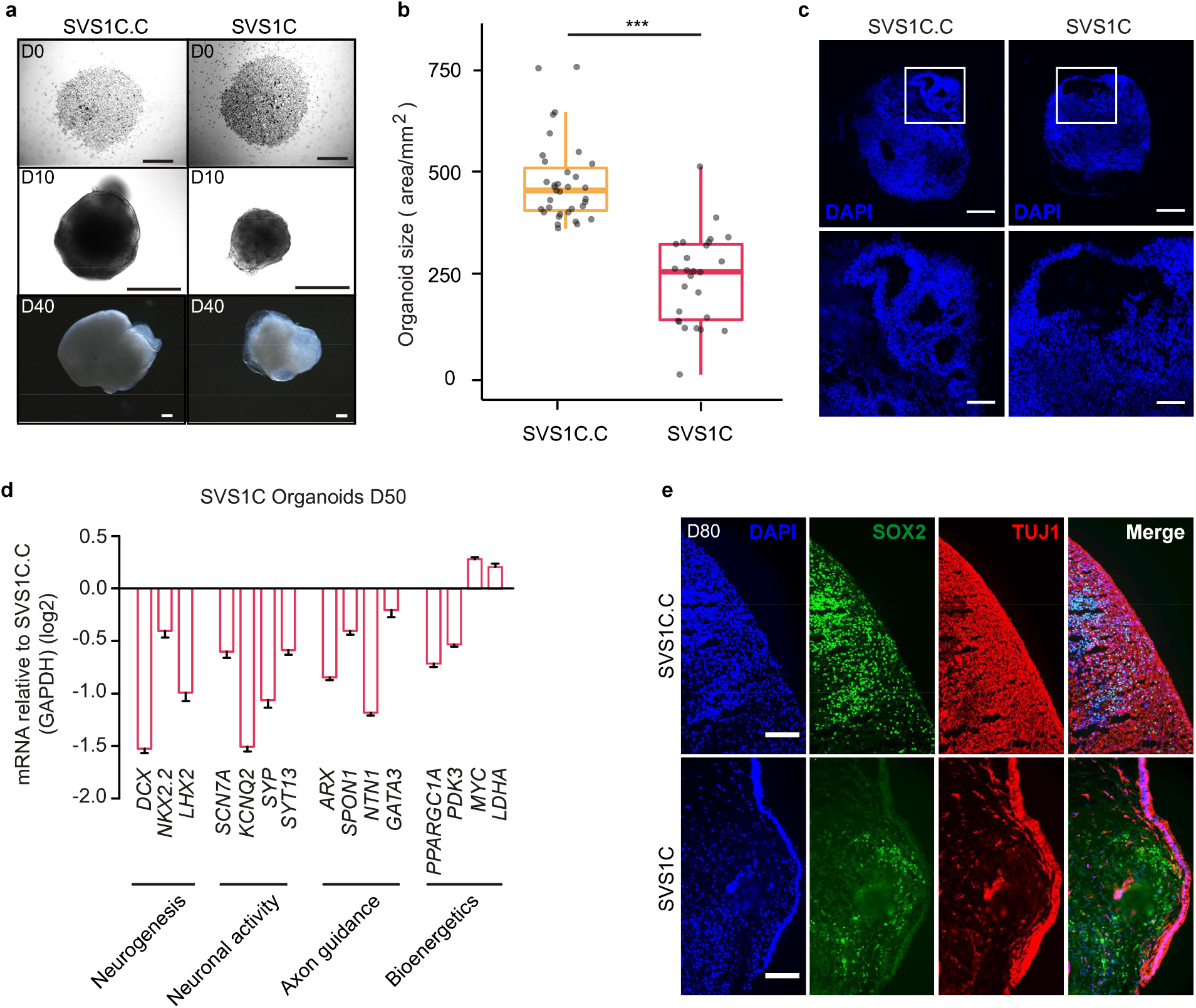
*SURF1* mutations impairs neurogenesis in iPSC-derived cerebral organoids. **a,** Representative images of cerebral organoids derived from SVS1C.C and SVS1C at various days of differentiation starting from the day in which organoids were embedded into Matrigel (day 0). Scale bars: DO = 500 μm, D10 = 200 μm, D40 = 200 μm. **b**, Quantification of the si¾ of cerebral organoids derived from SVS1C.C and SVS1C (horizmtal bars indicate the median, boxes are defined by the first and third quartiles, whiskers span 1.5 x inter-quartile range; n = 3 independent experiments; *** p < 0.001; Mann-Whitney t/test). **c**, Representative images of cerebral organoids derived from SVS1C.C and SVS1C showing the disruption of the epithelial layers in SVS1C. Scale bars in upper panel = 500 μm, in lower panel = 1000 μm. **d**, Quantitative real-time RT-PCR in cerebral organoids derived from SVS1C.C and SVS1C. Data were normali¾d to *GAPDH* (mean +/− s.d.). **e**, Representative images of cerebral organoids derived from SVS1C.C and SVS1C showing the reduced presence of SOX2-positive progenitors and TUJl-positive neurons. Scale bar: 100 μm.

In agreement with a neurogenic defect, SVS1C organoids showed downregulated expression of genes involved in neurogenesis like *NKX2.2, CDX, LHX2*, and *GATA3*, of genes regulating axon guidance like *ARX, NTN1*, and *SPON1*, and of genes associated with neuronal firing and synaptic activity like *SYP, SYT3, KCNQ2*, and *SCN7A* (**Fig. 5d**). SVS1C organoids also showed reduced expression of genes regulating mitochondrial metabolism like *PGC1A* and *PDK4*, while the expression of genes associated with glycolysis such as *MYC* and *LDHA* increased in comparison to isogenic control SVS1C.C organoids (**Fig. 5d**). Overall, the transcriptional profile of brain organoids was in line with the transcriptional profile of the two-dimensional (2D) neuronal cultures (**Fig. 2d-e; Fig. S5b-e; Fig. S6a**).

We then investigated the microscopic cytoarchitecture of the organoids. *SURF1* mutations altered the 3D self-organizing capacity of the cells and led to disrupted neuronal layering and reduced cortical thickness (**Fig. 5e**). SVS1C organoids showed aberrant distribution of the NPC marker SOX2 (**Fig. 5e; Fig. S8f**) and of the neuronal marker TUJ1 (**Fig. 5e**), while the expression of astrocyte marker S100ß appeared mainly unaltered (**Fig. S8e**). Altogether, the 3D model recapitulated the defects observed in the monolayer cultures and confirmed that SURF1 mutations impair the neural progenitor state, thereby disrupting the process of neurogenesis.

### Targeting NPC function allows the identification of *SURF1* gene augmentation as potential therapeutic strategy

Given the possible importance of dysfunctional neurogenesis in the pathogenesis of LS^SURF1^, we asked whether targeting NPC function could be of therapeutic benefit. We and others have previously employed iPSC-derived NPCs for identifying treatment interventions in the context of neuropsychiatric and mitochondrial diseases^31,32^. In comparison to isogenic control SVS1C.C-NPCs, SURF1-mutant SVS1C-NPCs showed bioenergetics defects, including reduced oxygen consumption (**Fig. 6a**) and increased lactate production (**Fig. 6b**). Neuronal profiling of NPC cultures revealed that early neurons within SVS1C-NPC cultures exhibited smaller neurites (**Fig. 6c**) and less branching points (**Fig. 6d**) than early neurons within SVS1C.C-NPC cultures. Hence, *SURF1* mutations hampered the development of physiological neuronal complexity, in agreement with what we observed in maturing DA neuronal cultures.

**Fig. 6.**
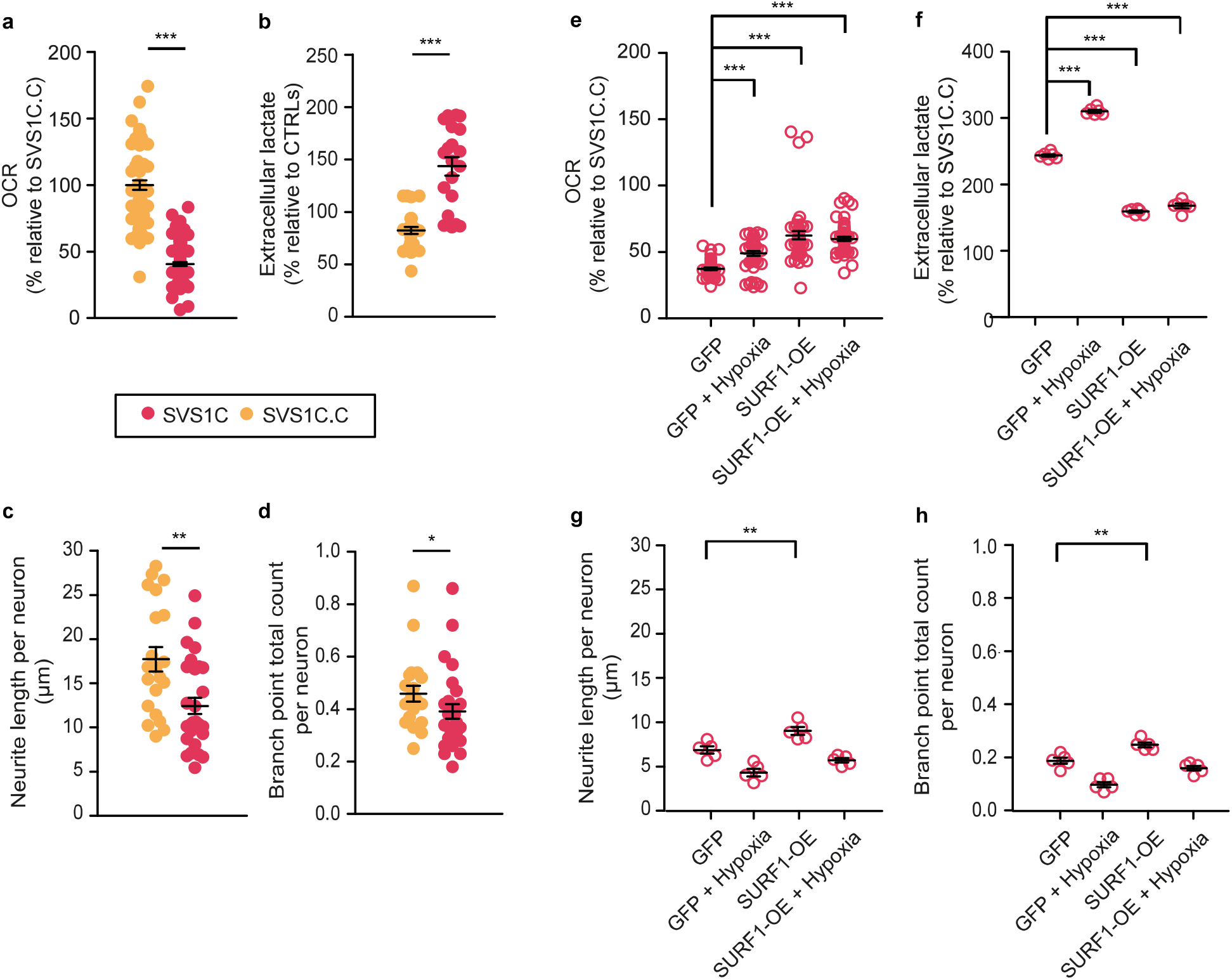
Restored functionality in *SURF1-*mutant NPCs upon wild-type SURF1 overexpression. **a,** OCR levels in NPCs from SVS1C.C and SVS1C (mean +/− s.e.m.; n = 3 independent experiments; *** p < 0.001; Mann-Whitney *U* test), **b**, Extracellular lactate in NPCs from SVS1C.C and SVS1C (mean +/− s.e.m.; n = 4 independent experiments; *** p < 0.001; Mann-Whitney *U* test), **c-d**, HCA-based quantification of neuronal branching in early neurons within NPC cultures from SVS1C.C and SVS1C (mean +/− s.e.m.; n = 3 independent experiments; * p < 0.05, ** p < 0.01; Mann-Whitney ř/test). **e**, OCR levels in SVSlC-NPCs stably expressing the GFP derivative mCitrine (GFP) or wild-type SURF1 (SURF1-OE) that were cultured overnight under normoxia or hypoxia (5% oxygen) (mean +/− s.e.m.; *** p < 0.001; one-way ANOVA followed by Bonferroni multiple comparison test), **g-h**, Neuronal profiling in SVSlC-NPCs expressing GFP or SURF1-OE that were cultured overnight under normoxic or hypoxic conditions (mean +/− s.e.m.; ** p < 0.01; one-way ANOVA followed by Bonferroni multiple comparison test).

We evaluated potential therapeutic strategies by measuring their ability to restore the functional and metabolic features of SURF1-NPCs. We first focused on interventions targeting OXPHOS activity. We tested: (i) the antioxidant α-Tocotrienol (AT3), whose *in* vivo-derived metabolite EPI-743 has been suggested as a therapy for LS^33^, (ii) fibroblast growth factor 21 (FGF21), which regulates glucose metabolism and promotes PGC1-α expression in human DA neurons^34,35^, (iii) glucose supplementation, is suggested for LS patients during acute metabolic decompensation^36^, and iv) pyruvate supplementation, which has been proposed as treatment for LS caused by COX defects^37^. We did not detect any consistent improvement of the phenotypes in NPCs exposed to these OXPHOS-centered interventions (**Fig. S9a-d**).

We then cultured NPCs under hypoxic conditions, which had shown functional benefit in animal models of LS with knock-out of the complex I gene *NDUFS4*^38^. Overnight culture under hypoxia (5% oxygen) significantly improved mitochondrial respiration in SURF1-NPCs (**Fig. 6e; Fig. S9i-j**). At the same time, however, hypoxia caused an undesired further increase in lactate production in SVS1C-NPCs (**Fig. 6f**), reduced OCR in control SVS1C.C-NPCs (**Fig. S9e-f**), and negatively affected neurogenesis in NPCs derived from both SVS1C (**Fig. 6g-h**) and SVS1C.C (**Fig. S9g-h**).

Next, we transduced SVS1C with a lentiviral vector encoding wild-type *SURF1* to assess the potential rescuing effect of SURF1 overexpression (SURF1-OE). In comparison to SVS1C transduced with lentiviruses encoding only the GFP derivative mCitrine (GFP), SURF1-OE showed significantly improved mitochondrial respiration (**Fig. 6e**) leading to a significant reduction of lactate production (**Fig. 6f**). Moreover, SURF1-OE improved the branching complexity of early neurons (**Fig. 6g-h**).

In conclusion, we used NPC function as a readout for interventions and found that SURF1-OE was beneficial in restoring bioenergetics and promoting neurogenesis. We thus propose *SURF1* gene augmentation therapy (GAT) as a potential strategy for restoring the neuronal function in LS^SURF1^ patients (**Fig. S9k**).

## Discussion

LS^SURF1^ has generally been considered an early-onset neurodegenerative disease. However, the mechanistic and pathophysiological understanding of LS^SURF1^ is hindered by the lack of effective model systems. *SURF1* knock-out has been carried out in several animals (flies, mice, zebrafish, and piglets) but all these efforts failed to fully recapitulate the neurological phenotypes of human LS^SURF1 14-18^. Here, we report the generation of a human mechanistic model of LS^SURF1^ based on patient-derived 2D neural cultures and 3D brain organoids. Using these models, we found that *SURF1* mutations impair the function and neurogenic potency of NPCs, ultimately leading to neurons with reduced complexity and activity.

Our data are in agreement with the clinical features of LS, which generally include developmental delay and cognitive impairment^6^. Microcephaly has often been reported in LS patients^2^.*SURF1* knock-out piglets, while not fully recapitulating the clinical phenotype of LS^SURF1^, also exhibited developmental alterations and reduced cortical thickness^18^. Moreover, a mutation in the mtDNA gene *MT-ATP6* that is associated with LS^39^ also caused abnormal NPC functionality^31^. Hence, it is possible that neurogenesis defects might contribute to other forms of LS and other mitochondrial disorders, which commonly involve severe neurodevelopmental manifestations^40^.

The dysfunction of SURF1-mutant NPCs might be related to their specific metabolic properties. Neurogenesis encompasses a series of complex cell fate transitions culminating in the terminal differentiation of NPCs into neurons. This process is accompanied by a metabolic shift from glycolysis to mitochondrial-based oxidative metabolism^41,42^. NPCs may represent the first cell type during this course to depend on mitochondrial oxidative function. Mitochondrial metabolism is in fact essential in mice for both embryonic and adult neurogenesis^43,44^. Moreover, we previously demonstrated that human iPSC-derived NPCs activate the OXPHOS machinery and are equipped with functionally mature mitochondria^31^. Should mitochondrial bioenergetics be impaired during brain development, NPCs might thus be the first cell population to be negatively affected. The dependence of cell type-specific function on different ATP-generating pathways might also explain the apparent absence of functional defects in astrocytes, given the fact that glial cells mainly rely on glycolytic metabolism^45,46^.

Our findings may open the way to new therapeutic avenues for LS focusing on improving neurogenesis rather than preventing the degeneration of fully functionally neurons. The current understanding that LS is caused by early-onset neurodegeneration had led to experimental treatment schemes centered around antioxidants to prevent the build-up of damaging free radicals^11,12^. However, blocking oxidative damage may not be sufficient in restoring neurogenesis. Antioxidant regiments may also dampen the physiological effect of radicals signaling, thereby blunting compensatory responses^47^. Other suggested treatment strategies aim at providing energy support through glucose or pyruvate administration^36,37^. But the need for bioenergetic restoration may not be uniform across all OXPHOS-dependent cell types. In LS^SURF1^, for example, muscle cells do not usually show the same level of impairment as neuronal cells^2^. Conversely, mutations in the gene encoding SCO2, which is a different assembly factor of COX, more strongly impair cardiac bioenergetics leading to cardiomyopathy rather than LS^48^. Hence, instead of general metabolic interventions, we propose that therapeutic strategies for LS^SURF1^ should specifically focus on improving neuronal metabolism.

Using NPC function as an interventional target, we discovered that overexpression of wild-type SURF1 restored neurogenesis in SURF1-mutant cells. There are ongoing clinical trials on gene augmentation therapy (GAT) for CNS disorders^49^. GAT also ameliorated the LS-like phenotypes in the complex I gene *NDUFS4* knock-out mice^50^. Given the proposed impairment of neurogenesis, we think that GAT in LS^SURF1^ should be carried out as early as possible to enable the formation of functional neurons and neuronal networks in the patients. Further studies are needed to investigate in details the potential adverse reactions of *SURF1*-GAT in live animals and to identify the most effective strategy with respect to viral vector types, timing, and sites of delivery.

In conclusion, we generated the first human mechanistic models of LS^SURF1^, which enabled us to discover that SURF1 defects impair neurogenesis and led us to propose early-initiated *SURF1*-GAT as a potential therapeutic strategy for counteracting the neurodevelopmental failure in this fatal neuropediatric disorder.

## Supporting information

Supp Figures and Full Methods

## Author contributions

Conceptualization, A.P., G.I., M.S. Methodology, G.I., A.R-W., P.L., R.J., A.Z., C.S., W.S., T.H., U.H.C., S.D.; Formal Analysis, B.M., P.G.; Resources, E.E.W; W.S.; N.R.; R.K., M.S.; Writing –Original Draft, A.P.; Writing – Review & Editing, A.P., M.S., J.P., E.E.W.; Supervision, A.P., M.G., R.K., E.E.W., N.R., J.P.; Visualization, G.I., A.P.; Funding Acquisition, A.P., M.S.

## Acknowledgements

We thank Carlo Viscomi (MRC-MBU, Cambridge UK) for critical reading of the manuscript and Carmen Lorenz (MDC) for proofreading. We are grateful to Anje Sporbert (Advanced Light Microscopy, MDC) for help with the mitochondrial movement recordings and to the Viral Core Facility of the Charité for generating lentiviral particles. We are thankful to Celina Kassner, Pia Badstuebner, and Luica Delius for technical support with cell culture experiments. We acknowledge support from the MDC (BOOST award to A.P. and M.S.), the Deutsche Forschungsgemeinschaft (DFG) (PR1527/1-1 & SCHU1187/4-1 to A.P. and M.S.; Neurocure Exc 257 to M.S.), the Berlin Institute of Health (BIH) (BIH Validation Funds to A.P. and M.S.), the National Science Center of Poland (No. 2016/22/M/NZ2/00548 to P.L), and the Bundesministerium für Bildung und Forschung (BMBF) (e:Bio young investigator grant AZ.031A318 to A.P. and AERIAL P1 to J.P.).

