## Supplementary material for "*SURF1* mutations causative of Leigh syndrome impair human neurogenesis": Supp Figures and Full Methods

Figure S1

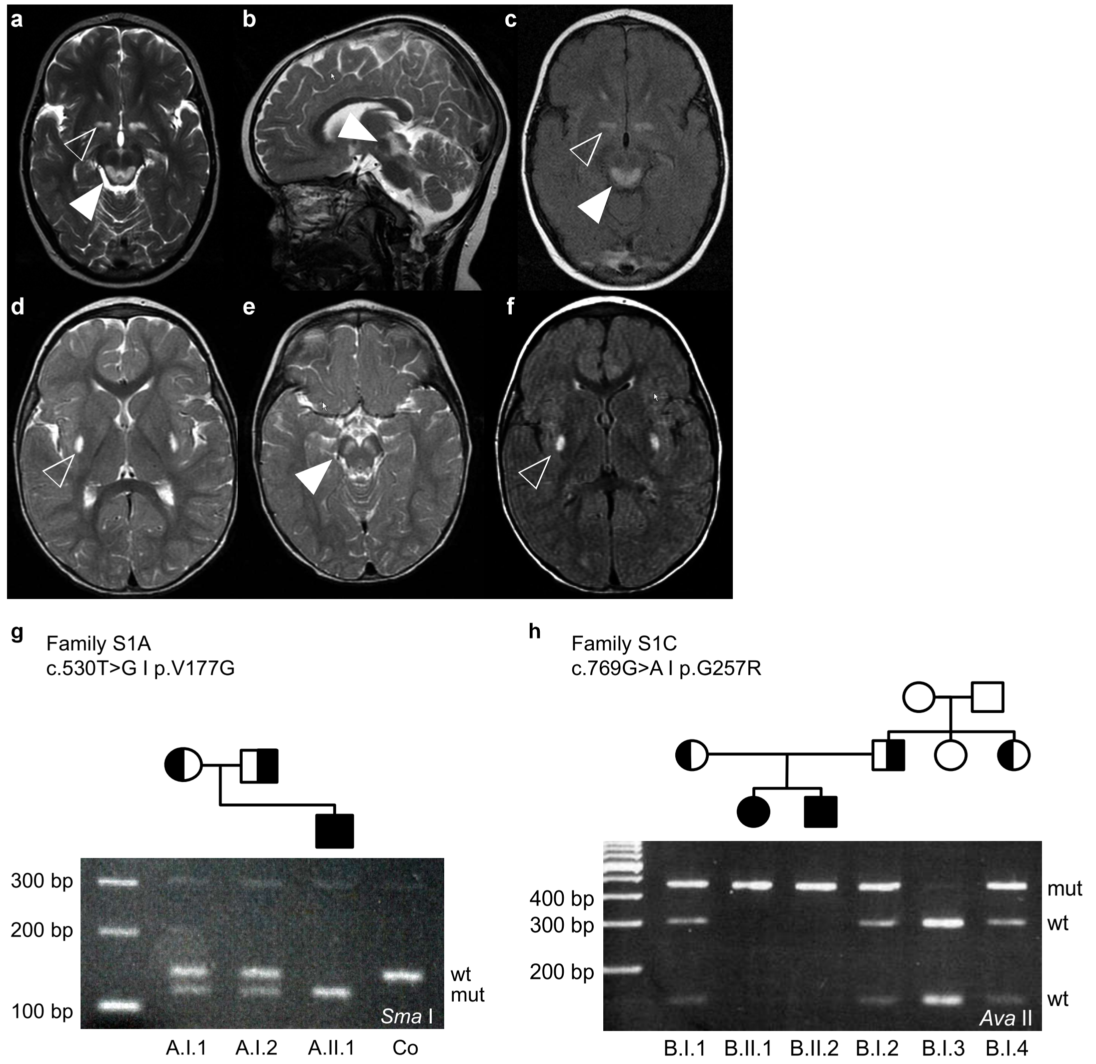
a
b
c
d
e
f
g
Family S1A
c.530T>G I p.V177G
300 bp
200 bp
100 bp
wt
mut
Sma I
A.I.1
A.I.2
A.II.1
Co
h
Family S1C
c.769G>A I p.G257R
400 bp
300 bp
200 bp
mut
wt
wt
Ava II
B.I.1
B.II.1
B.II.2
B.I.2
B.I.3
B.I.4

**Fig. S1. Clinical features of LS$^{SURF1}$ patients (relative to Fig. 1). a-f,** Cranial MRI images of S1A patient (a-c) and of S1C patient (d-e). Open arrowheads depict $T_2$-signal intense lesions in the basal ganglia as the pathological hallmark of LS. Closed arrowheads point at brainstem lesions comprising the *formatio reticularis* (a-c) and the *substantia nigra* (e). **g-h**, Pedigrees of the families and segregation between the LS phenotype with homozygosity for the *SURF1* mutation in the families of the S1A patient (A.II.1) and S1C patient (B.II.2). The mutations c.530T>G and c.769G>A were visualized *via* primer induced restriction analysis (PIRA) using *SmaI* and *AvaII*, respectively

Figure S2

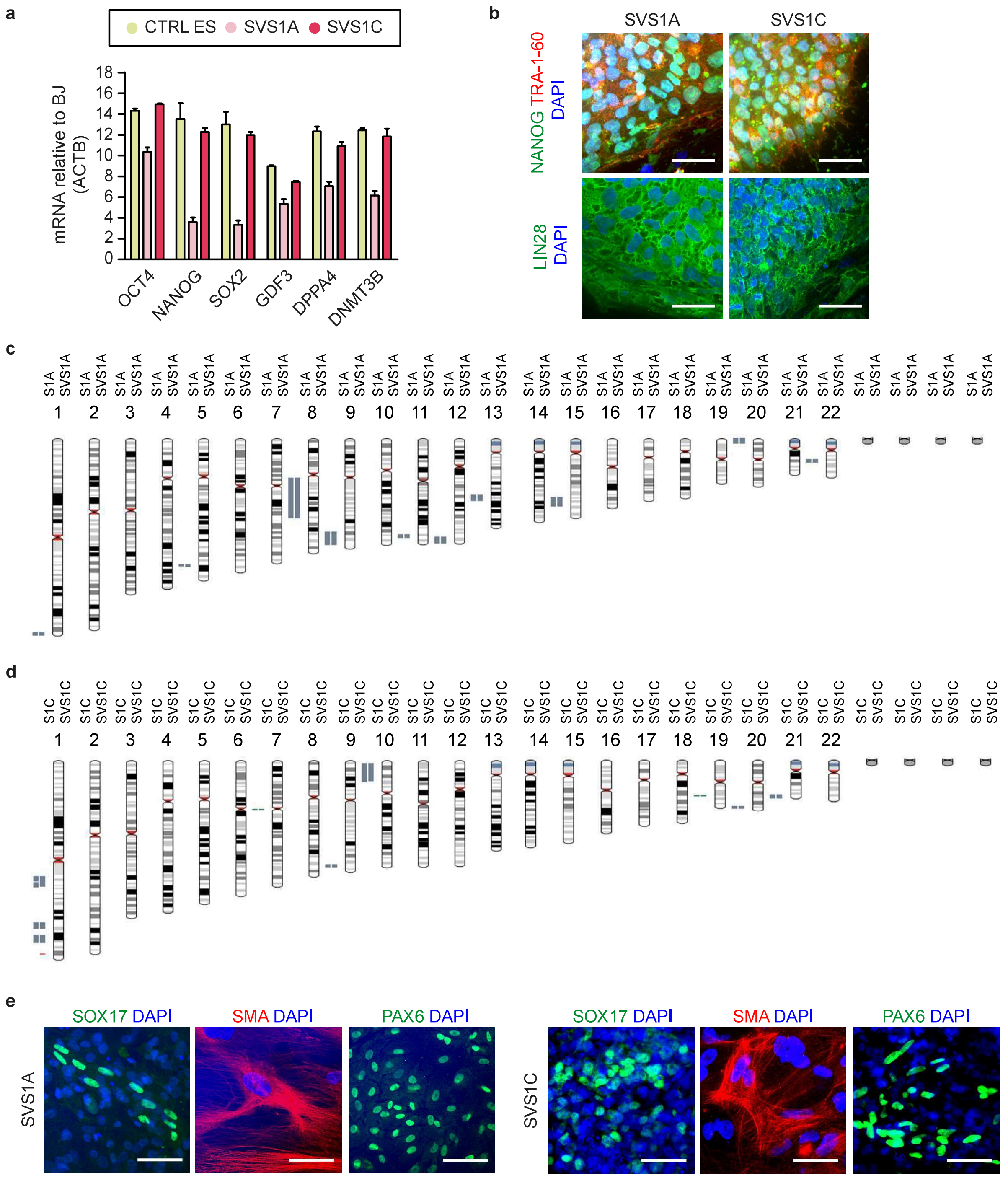
a
CTRL ES SVS1A SVS1C
mRNA relative to BJ (ACTB)
0 2 4 6 8 10 12 14 16 18
OCT4 NANOG SOX2 GDF3 DPPA4 DNMT3B
b
SVS1A SVS1C
NANOG TRA-1-60 DAPI
LIN28 DAPI
c
S1A SVS1A
1 2 3 4 5 6 7 8 9 10 11 12 13 14 15 16 17 18 19 20 21 22
d
S1C SVS1C
1 2 3 4 5 6 7 8 9 10 11 12 13 14 15 16 17 18 19 20 21 22
e
SOX17 DAPI
SMA DAPI
PAX6 DAPI
SVS1A
SVS1C

**Fig. S2. Characterization of iPSCs from LS$^{SURF1}$ patients (SURF1 iPS) (relative to Fig. 1). a,** Quantitative real-time RT-PCR of pluripotency-associated markers in CTRL ES (H1) and SURF1 iPS (SVS1A and SVS1C). Data were normalized to *ACTB* (mean +/- s.d.; n = 2 independent experiments). **b**, Representative images of pluripotency-associated marker proteins in SVS1A and SVS1C. Scale bar: 50 µm. **c-d**, Karyotype analysis confirmed that SVS1A and SVS1C carried a normal karyotype and were derived from the original fibroblasts S1A and S1C, respectively. **e**, SVS1A and SVS1C showed pluripotency features, exemplified here with the ability to give rise to cells belonging to the three germ layers: endoderm (SOX17), mesoderm (smooth muscle actin, SMA), and ectoderm (PAX6). Scale bar: 50 µm.

### Figure S3

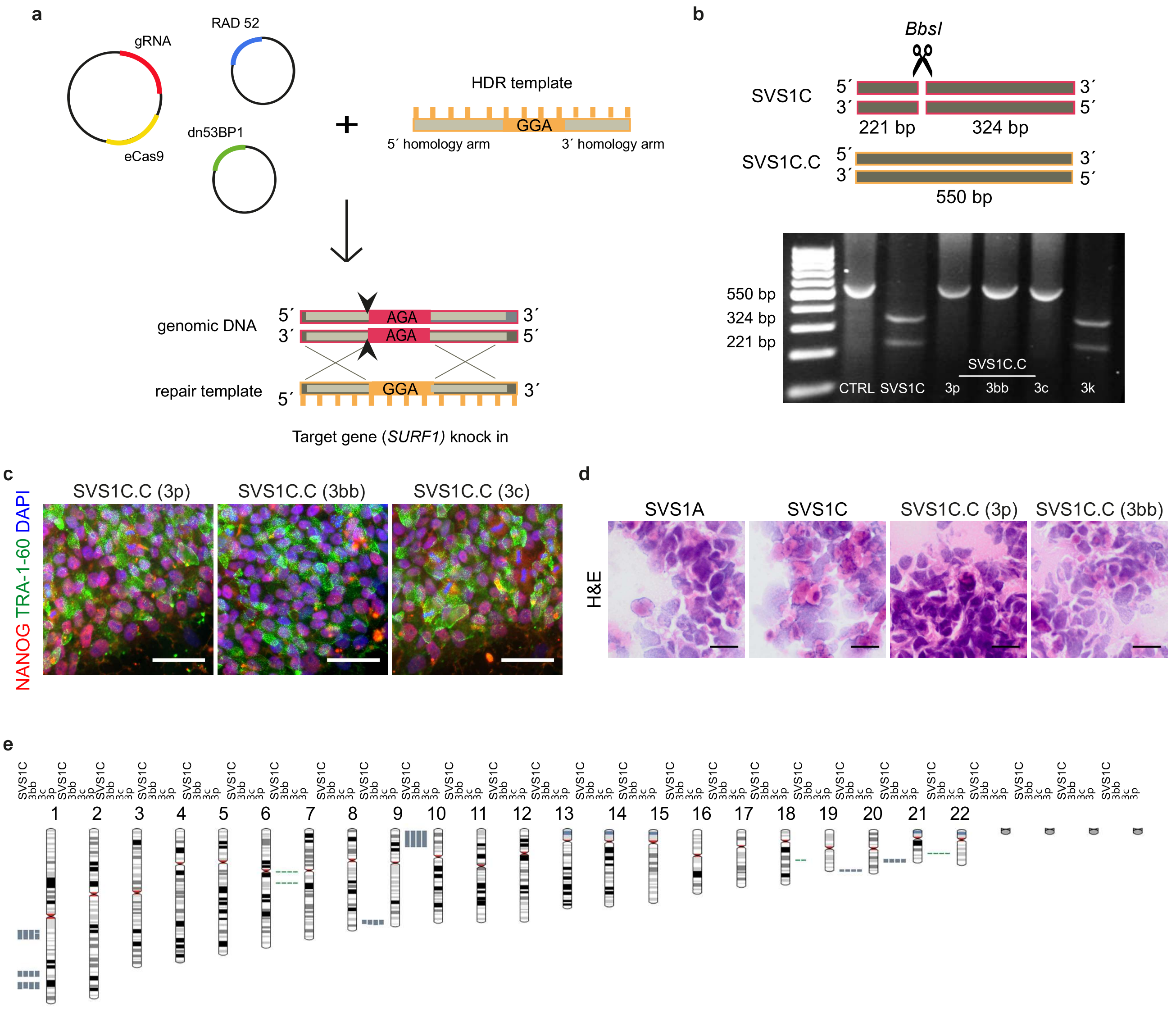

a
gRNA
RAD 52
eCas9
dn53BP1
HDR template
GGA
5´ homology arm
3´ homology arm
genomic DNA
AGA
repair template
Target gene (SURF1) knock in
b
BbsI
SVS1C
221 bp
324 bp
SVS1C.C
550 bp
550 bp
324 bp
221 bp
CTRL SVS1C 3p 3bb 3c 3k
c
NANOG TRA-1-60 DAPI
SVS1C.C (3p)
SVS1C.C (3bb)
SVS1C.C (3c)
d
H&E
SVS1A
SVS1C
SVS1C.C (3p)
SVS1C.C (3bb)
e
1 2 3 4 5 6 7 8 9 10 11 12 13 14 15 16 17 18 19 20 21 22

**Fig. S3. Genome editing of SURF1 iPS (relative to Fig. 1). a,** Schematic of the CRISPR/Cas9 knock-in strategy showing the plasmids and the homologous direct repair (HDR) template used. **b**, To confirm the presence of the c.769G>A mutation, we used primer induced restriction analysis (PIRA). The PCR product of 550 bp was cut by *BbsI* into 221+324 bp fragments only in the presence of the mutation c.769G>A. **c**, Representative images of pluripotency-associated markers NANOG and TRA-1-60 in SVS1C.C (clones 3p, 3bb, and 3c). Scale bar: 50 μm. **d**, Hematoxylin eosin (H&E) staining of the images reported in Fig. 1f. Scale bar: 50 μm. **e**, Karyotype analysis confirmed that the three SVS1C.C clones (3bb, 3c, and 3p) had a normal karyotype and that were derived from the original iPSC line SVS1C.

Figure S4

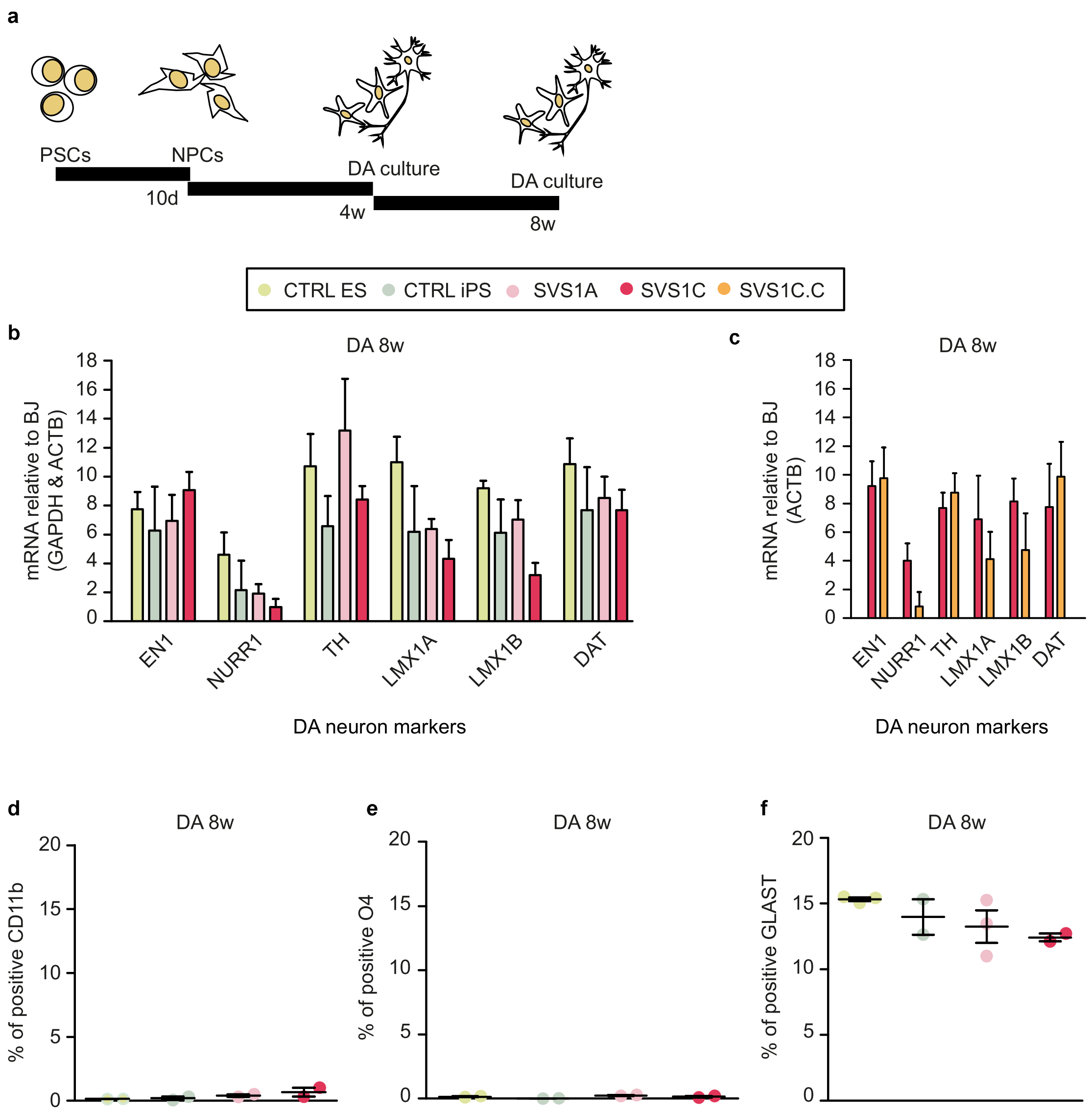

a
PSCs
NPCs
DA culture
DA culture
10d
4w
8w
CTRL ES
CTRL iPS
SVS1A
SVS1C
SVS1C.C
b
DA 8w
mRNA relative to BJ (GAPDH & ACTB)
EN1
NURR1
TH
LMX1A
LMX1B
DAT
DA neuron markers
c
DA 8w
mRNA relative to BJ (ACTB)
EN1
NURR1
TH
LMX1A
LMX1B
DAT
DA neuron markers
d
DA 8w
% of positive CD11b
e
DA 8w
% of positive O4
f
DA 8w
% of positive GLAST

**Fig. S4. Characterization of dopaminergic-enriched (DA) neuronal cultures (relative to Fig. 2). a,** Schematic of the differentiation from pluripotent stem cells (PSCs) to neural progenitor cells (NPCs) and to dopaminergic (DA) neuronal cultures. **b**, Quantitative real-time RT-PCR of genes associated with DA neuronal identity in 8w DA cultures from CTRL ES (H1), CTRL iPS (XM001 and TFBJ), SVS1A, and SVS1C. Data were normalized to both *GAPDH* and *ACTB* (mean +/- s.d.; n = 4 independent experiments). **c**, Quantitative real-time RT-PCR of DA markers in 8w DA cultures from SVS1C.C and SVS1C. Data were normalized to *ACTB* (mean +/- s.d.; n = 3 independent experiments). **d-f,** MACs-based quantification of glial surface markers: CD11b for microglia, O4 for oligodendrocytes, and GLAST for astrocytes in CTRL ES (H1), CTRL iPS (XM001) and SURF1 iPS (SVS1A and SVS1C). (mean +/- s.e.m.; n = 2 independent experiments; one-way ANOVA followed by Bonferroni multiple comparison test; not significant).

Figure S5

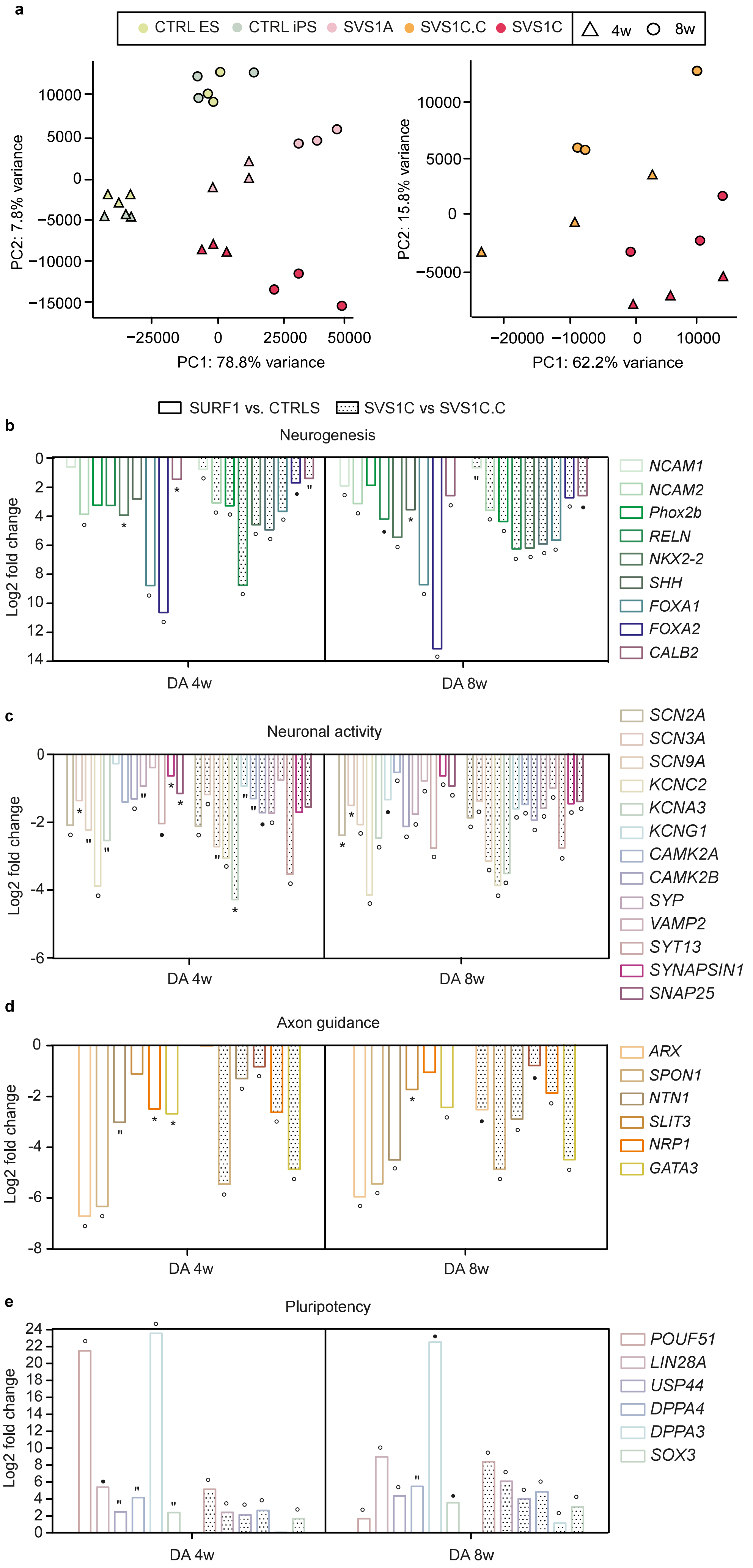

a
CTRL ES
CTRL iPS
SVS1A
SVS1C.C
SVS1C
4w
8w
PC2: 7.8% variance
PC1: 78.8% variance
PC2: 15.8% variance
PC1: 62.2% variance
SURF1 vs. CTRLS
SVS1C vs SVS1C.C
b
Neurogenesis
Log2 fold change
DA 4w
DA 8w
NCAM1
NCAM2
Phox2b
RELN
NKX2-2
SHH
FOXA1
FOXA2
CALB2
c
Neuronal activity
SCN2A
SCN3A
SCN9A
KCNC2
KCNA3
KCNG1
CAMK2A
CAMK2B
SYP
VAMP2
SYT13
SYNAPSIN1
SNAP25
d
Axon guidance
ARX
SPON1
NTN1
SLIT3
NRP1
GATA3
e
Pluripotency
POUF51
LIN28A
USP44
DPPA4
DPPA3
SOX3

**Fig. S5. Transcriptome analysis of DA neuronal cultures from LS$^{SURF1}$ patients (relative to Fig. 2). a,** Left panel: principal component analysis (PCA) of mRNA-based transcriptome (polyA) of 4w and 8w DA cultures derived from SURF1 iPS (SVS1A and SVS1C) *vs.* CTRLS (H1 and XM001) (n = 3 independent experiments). Right panel: PCA of total RNA-based transcriptome of 4w and 8w DA cultures derived from SVS1C *vs.* SVS1C.C (n = 3 independent experiments). **b-e**, LFC of genes regulating neurogenesis, neuronal activity, axon guidance, and pluripotency in 4w and 8w DA cultures from SURF1 iPS (SVS1A and SVS1C) *vs.* CTRL (H1 and XM001) (empty bars) and in 4w and 8w DA cultures from SVS1C *vs.* SVS1C.C (dotted bars) (* $p < 0.05$, ′′ $p < 0.01$, o $p < 0.001$, ● $p < 0.0001$).

### Figure S6

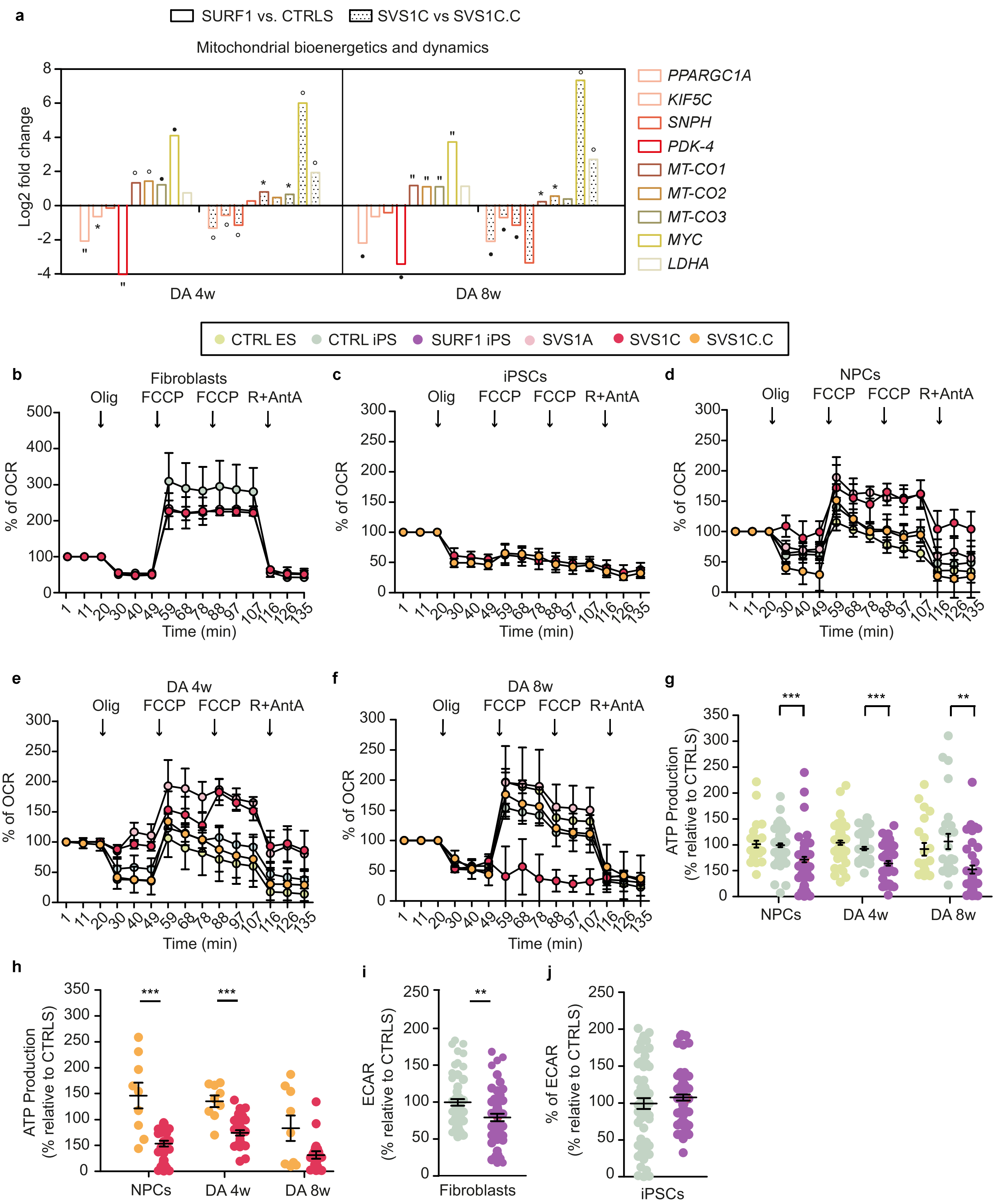
a
SURF1 vs. CTRLS
SVS1C vs SVS1C.C
Mitochondrial bioenergetics and dynamics
Log2 fold change
DA 4w
DA 8w
PPARGC1A
KIF5C
SNPH
PDK-4
MT-CO1
MT-CO2
MT-CO3
MYC
LDHA
CTRL ES
CTRL iPS
SURF1 iPS
SVS1A
SVS1C
SVS1C.C
b
Fibroblasts
c
iPSCs
d
NPCs
e
DA 4w
f
DA 8w
Olig
FCCP
R+AntA
% of OCR
Time (min)
g
ATP Production
(% relative to CTRLS)
NPCs
h
i
ECAR
Fibroblasts
j
% of ECAR

**Fig. S6. Bioenergetic profiling of NPCs and DA cultures from $LS^{SURF1}$ patients (relative to Fig. 2 and Fig.3). a**, LFC of genes regulating mitochondrial bioenergetics and dynamics in 4w and 8w DA cultures from SURF1 iPS (SVS1A and SVS1C) *vs.* CTRLS (H1 and XM001) (empty bars) and in 4w and 8w DA cultures from SVS1C *vs.* SVS1C.C (dotted bars) (* $p < 0.05$, ´´ $p < 0.01$, o $p < 0.001$, ● $p < 0.0001$). **b**, OCR profile in fibroblasts derived from $LS^{SURF1}$ patients (S1A and S1C) and in fibroblasts from healthy controls (CON1 and BJ) (mean +/- s.e.m.; n = 2 independent experiments). **c**, OCR profile in SVS1C and SVS1C.C (mean +/- s.e.m.; n = 2 independent experiments). **d**, OCR profile in NPCs from CTRL ES (H1), CTRL iPS (XM001 and TFBJ), SURF1 iPS (SVS1A and SVS1C), and SVS1C.C (mean +/- s.e.m.; n = 2 independent experiments). **e-f**, OCR profile in 4w and 8w DA cultures derived from CTRL ES (H1), CTRL iPS (XM001), SURF1 iPS (SVS1A and SVS1C), and SVS1C.C (mean +/- s.e.m.; n = 2 independent experiments). **g**, ATP production based on OCR consumption following oligomycin treatment in NPCs, 4w, and 8w DA cultures from CTRL ES (H1), CTRL iPS (XM001), and SURF1 iPS (SVS1A and SVS1C) (mean +/- s.e.m.; n = 3 independent experiments; ** $p < 0.01$, *** $p < 0.001$; one-way ANOVA followed by Bonferroni multiple comparison test). **h**, ATP production in NPCs, 4w, and 8w DA cultures from SVS1C.C and SVS1C (mean +/- s.e.m.; *** $p < 0.001$; Mann-Whitney *U* test). **i**, Extracellular acidification rate (ECAR) in fibroblasts derived from $LS^{SURF1}$ patients (S1A and S1C) and in fibroblasts from healthy controls (CON1 and BJ) (mean +/- s.e.m.; n = 2 independent experiments; ** $p < 0.01$; Mann-Whitney *U* test). **j**, ECAR in CTRL iPS (XM001) and SURF1 iPS (SVS1A and SVS1C) (mean +/- s.e.m.; n = 2 independent experiments; Mann-Whitney *U* test; not significant).

### Figure S7

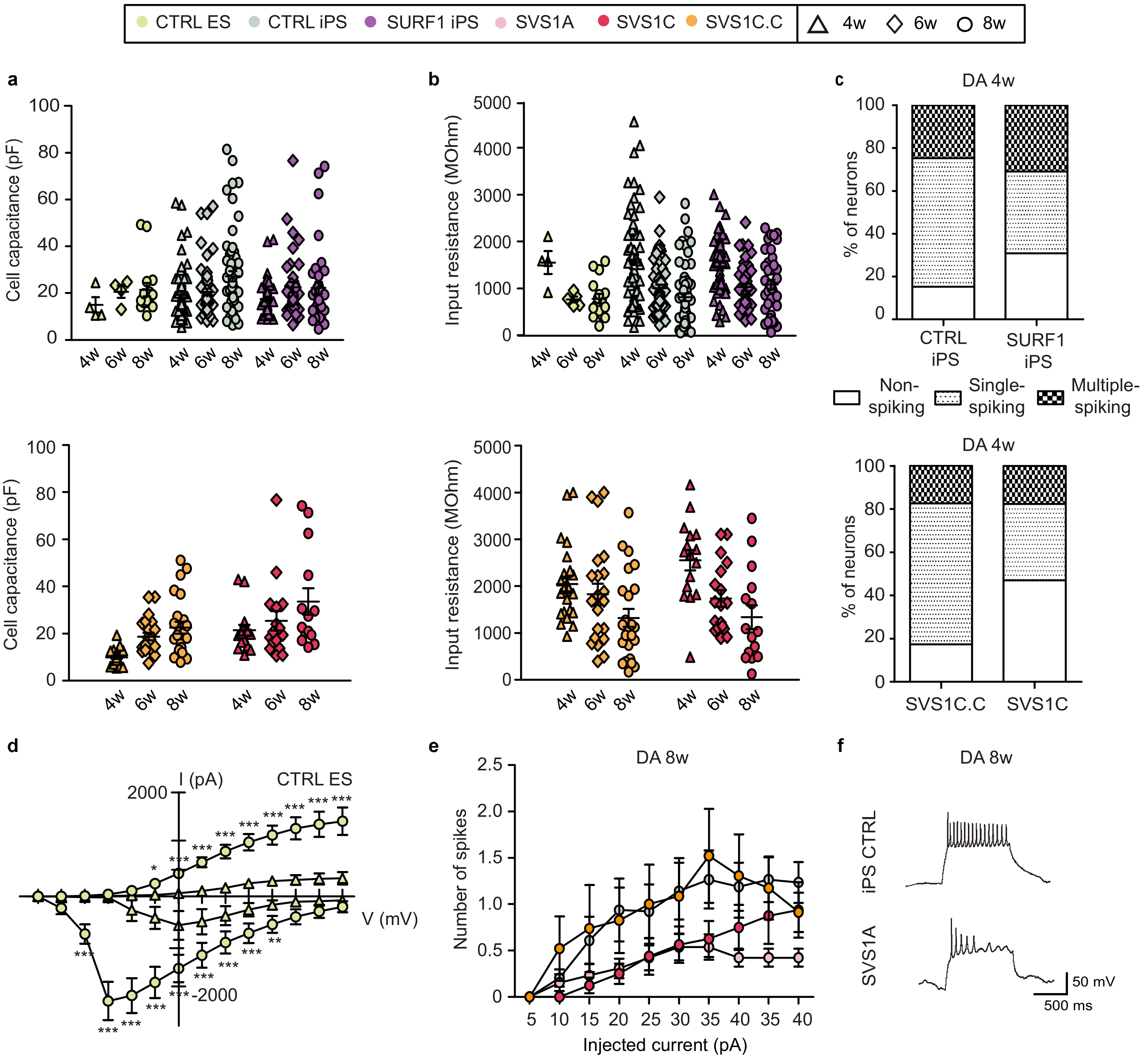
CTRL ES
CTRL iPS
SURF1 iPS
SVS1A
SVS1C
SVS1C.C
4w
6w
8w
a
Cell capacitance (pF)
b
Input resistance (MOhm)
c
DA 4w
% of neurons
CTRL iPS
SURF1 iPS
Non-spiking
Single-spiking
Multiple-spiking
SVS1C.C
SVS1C
d
I (pA)
CTRL ES
V (mV)
e
DA 8w
Number of spikes
Injected current (pA)
f
DA 8w
iPS CTRL
SVS1A
50 mV
500 ms

**Fig. S7. Electrophysiological properties of DA cultures from LS$^{SURF1}$ patients (relative to Fig. 4). a**, Cell capacitance (in picofarad, pF) in DA neuronal cultures from CTRL ES (H1), CTRL iPS (XM001 and TFBJ), and SURF1 iPS (SVS1A and SVS1C) (above) and in SVS1C and SVS1C.C (below) (mean +/- s.d.; at least n = 2 independent experiments). **b**, Input resistance (in MilliOhm, MOhm) in DA neuronal cultures from CTRL ES (H1), CTRL iPS (XM001), and SURF1 iPS (SVS1A and SVS1C) (above) and in SVS1C and SVS1C.C (below) (mean +/- s.d.; at least n = 2 independent experiments). **c**, Percentage of neurons (mean) that are not-spiking, single-spiking, or multiple spiking in 4w DA cultures from CTRL iPS (XM001 and TFBJ) and SURF1 iPS (SVS1A and SVS1C) (above) and from SVS1C.C and SVS1C (below) (at least n = 2 independent experiments). **d**, Sodium and potassium currents in 4w and 8w DA cultures from CTRL ES (H1) (mean +/- s.d.; n=3 independent experiments; * $p < 0.05$, ** $p < 0.01$, *** $p < 0.001$; two-way ANOVA). **e**, Number of spikes in 8w DA cultures from CTRL iPS (XM001 and TFBJ), SURF1 iPS (SVS1A and SVS1C) (above), and SVS1C.C (below) (mean +/- s.e.m.; at least n = 2 independent experiments). **f**, Representative electrophysiology traces showing that 8w DA cultures from SVS1A contained neurons with less multiple spiking activity compared to neurons from CTRL iPS (XM001).

Figure S8

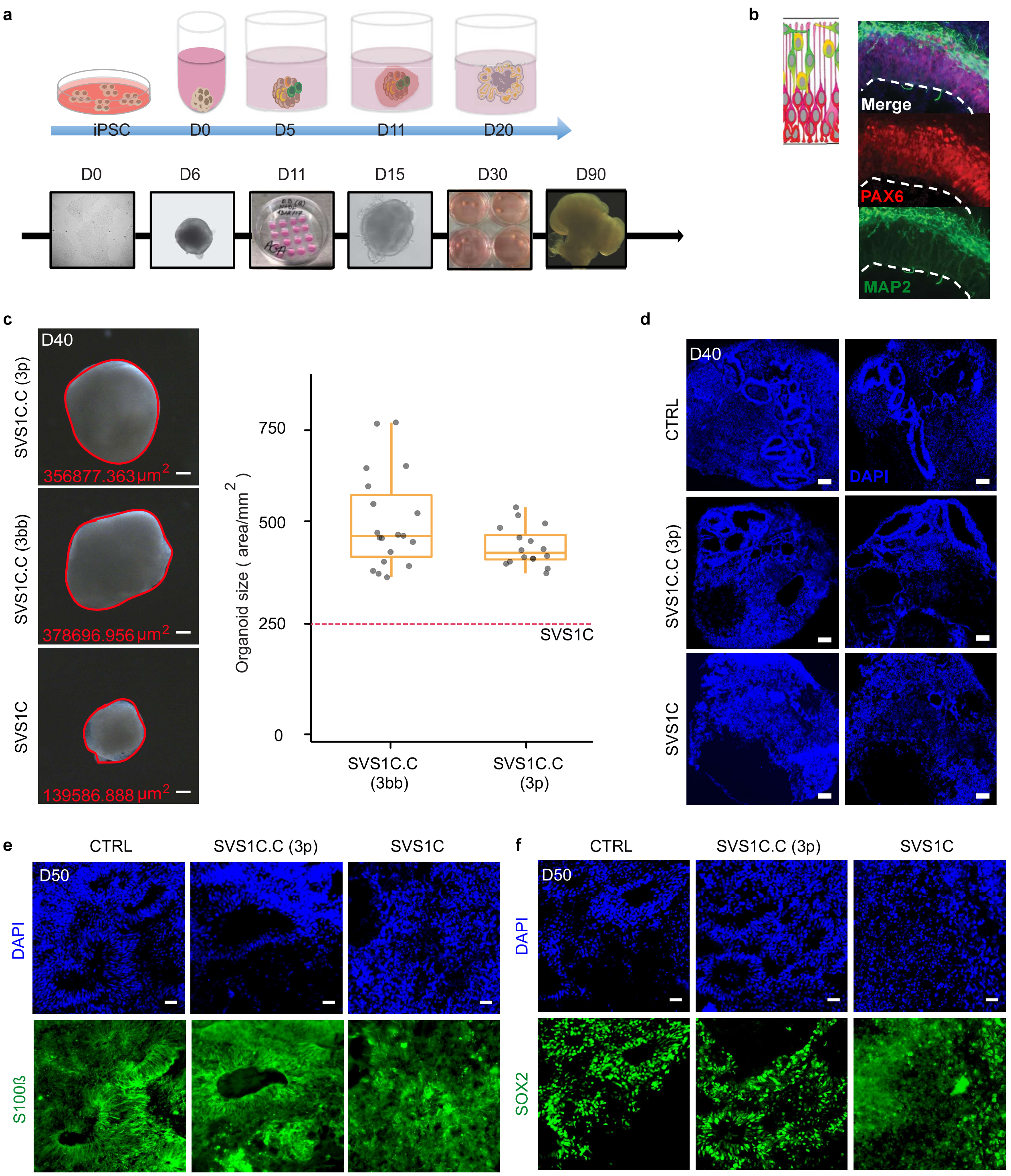

a
iPSC
D0
D5
D11
D20
D0
D6
D11
D15
D30
D90
b
Merge
PAX6
MAP2
c
D40
SVS1C.C (3p)
356877.363 µm2
SVS1C.C (3bb)
378696.956 µm2
SVS1C
139586.888 µm2
Organoid size ( area/mm2 )
750
500
250
0
SVS1C
SVS1C.C (3bb)
SVS1C.C (3p)
d
D40
CTRL
DAPI
SVS1C.C (3p)
SVS1C
e
CTRL
SVS1C.C (3p)
SVS1C
D50
DAPI
S100ß
f
CTRL
SVS1C.C (3p)
SVS1C
D50
DAPI
SOX2

**Fig. S8. Characterization of iPSC-derived cerebral organoids from LS$^{SURF1}$ patients (relative to Fig. 5). a**, Schematic of the generation of cerebral organoids from iPSCs. **b**, Representative images from control brain organoids showing the distribution of progenitors and neuronal cells in the self-organized epithelial layers. **c**, Left panel: representative macroscopic images of brain organoids at day 40. Scale bar: 100 µm. Right panel: quantification of the size of cerebral organoids demonstrated that the two clones of SVS1C.C (3p and 3bb) had similar increase in size as compared to organoids from SVS1C (red horizontal line) (horizontal bars indicate the median, boxes are defined by the first and third quartiles, whiskers span 1.5 x inter-quartile range; n = 2 independent experiments). **d**, Representative images depicting the epithelial layers present in organoids from CTRL iPS (XM001) that were disrupted in SVS1C organoids but were restored in organoids from SVS1C.C. Scale bar: 100 µm. **e-f**, Representative images of astrocytes (S100β-positive) and NPCs (SOX2-positive) in organoids from CTRL iPS (XM001), SVS1C.C, and SVS1C. Scale bar: 100 µm.

### Figure S9

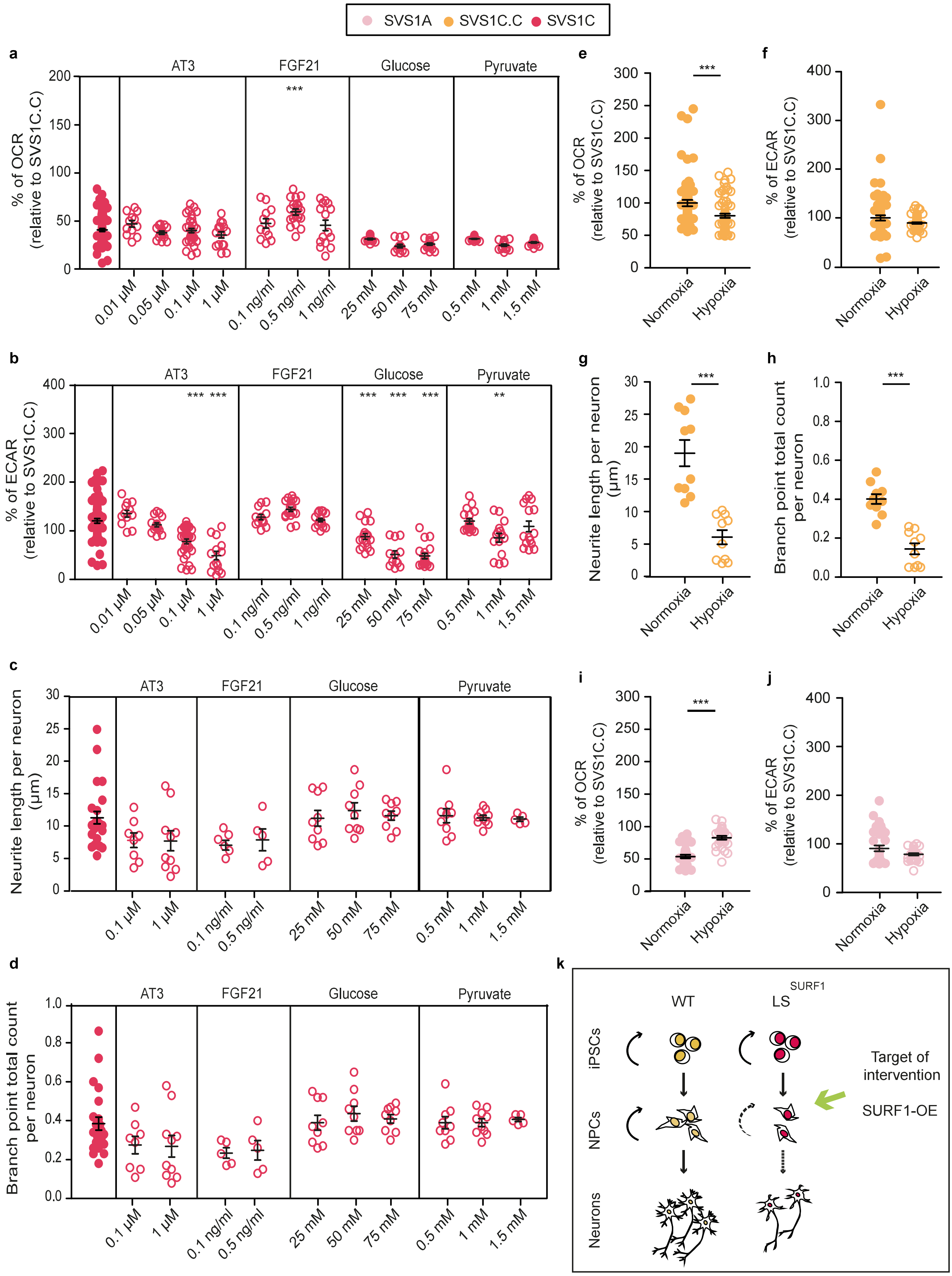

SVS1A
SVS1C.C
SVS1C
a
% of OCR (relative to SVS1C.C)
AT3
FGF21
Glucose
Pyruvate
0.01 µM
0.05 µM
0.1 µM
1 µM
0.1 ng/ml
0.5 ng/ml
1 ng/ml
25 mM
50 mM
75 mM
0.5 mM
1 mM
1.5 mM
b
% of ECAR (relative to SVS1C.C)
c
Neurite length per neuron (µm)
d
Branch point total count per neuron
e
f
g
h
i
j
Normoxia
Hypoxia
k
WT
LS SURF1
iPSCs
NPCs
Neurons
Target of intervention
SURF1-OE

**Fig. S9. Treatment strategies in NPCs from LS$^{SURF1}$ patients (relative to Fig. 6). a-b,** OCR and ECAR levels in NPCs from SVS1C under untreated conditions and following overnight treatment with α-Tocotrienol (AT3), FGF21, glucose, or pyruvate (mean +/- s.e.m.; ** $p < 0.01$, *** $p < 0.001$; one-way ANOVA followed by Bonferroni multiple comparison test). **c-d**, Neuronal profiling in SVS1C-NPCs under untreated conditions and following overnight treatment with AT3, FGF21, glucose, or pyruvate (mean +/- s.e.m.; one-way ANOVA followed by Bonferroni multiple comparison test; not significant). **e**, OCR levels in SVS1C.C-NPCs cultured overnight under normoxia or hypoxia (5% oxygen) (mean +/- s.e.m.; *** $p < 0.001$; Mann-Whitney *U* test). **f**, ECAR levels in SVS1C.C-NPCs cultured overnight under normoxia or hypoxia (mean +/- s.e.m.; Mann-Whitney *U* test; not significant). **g-h**, Neuronal profiling in SVS1C.C-NPCs cultured overnight under normoxia or hypoxia (mean +/- s.e.m.; *** $p < 0.001$; Mann-Whitney *U* test). **i**, OCR levels in SVS1A-NPCs cultured overnight under normoxia or hypoxia (mean +/- s.e.m.; *** $p < 0.001$; Mann-Whitney *U* test). **j**, ECAR levels in SVS1A-NPCs cultured overnight under normoxia or hypoxia (mean +/- s.e.m.; Mann-Whitney *U* test, treated vs. untreated; not significant). **k**, Schematic depicting the suggested mechanisms caused by *SURF1* mutations involving NPC defects that can be exploited to identify treatment strategies such as the one here proposed based on SURF1-OE.

#### Full methods

##### Subject details

Written informed consent to use patient material was obtained from the guardians according to the Declaration of Helsinki. The study was approved by the IRB of the Charité (EA2/131/13 and EA2/107/14). Patient fibroblasts S1A and S1C were obtained from two distinct consanguineous families **(Fig. S1)**. S1A patient was male with healthy consanguineous parents. Diagnosis of $LS^{SURF1}$ included cranial MRI showing bilateral basal ganglia necroses and $T_2$-signal intensities in the brainstem, elevated lactate in the cerebrospinal fluid (CSF, 4.3 mmol/l; N: <2), pronounced reduction of COX staining and COX activity in muscle (15 mU/mg NCP; N: 90-281) and in cultured skin fibroblasts (130 mU/U CS; N: 680-1190), with normal enzyme activities of the other respiratory chain complexes. Sequencing of *SURF1* showed a homozygous variant (c.530T>G, NM_003172) in exon 6 that was heterozygous in both parents. The skin biopsy was taken at 9 years of age. Patient S1A died at 25 years of age during a metabolic crisis in the course of an infection. Patient S1C was male with healthy consanguineous parents. Cranial MRI revealed the characteristic basal ganglia necroses of LS and biochemical investigation confirmed COX deficiency in muscle (158 mU/U CS; N: 520-2080) and in cultured fibroblasts (100 mU/U CS; N: 342-627), with normal activity of the other OXPHOS complexes. Sequence analysis of *SURF1* detected a homozygous c.769G>A variant in exon 8 causing the exchange of a highly conserved glycine for an arginine (p.G257R). The skin biopsy was taken at 20 months of age. Patient S1C died from global respiratory and cardiac failure at the age of 5 years.

##### iPSC generation

We reprogrammed patient fibroblasts using Sendai viruses (CytoTune-iPS 2.0 from Thermo Fisher Scientific). hESC line H1 was purchased from WiCell and was used in accordance to the German license of Alessandro Prigione issued by the Robert Koch Institute

(AZ: 3.04.02/0077-E01). Control iPSC lines were previously generated using episomal plasmids [1] and described as TFBJ[2] and XM001[3]. All pluripotent stem cells (PSCs) were cultivated on Matrigel (BD Bioscience)-coated plates using StemMACS iPS-Brew XF medium (Miltenyi Biotec GmbH, #130-104-368), supplemented with Pen/Strep (Thermo Fisher Scientific) and MycoZap (Lonza). We routinely monitored against mycoplasma contamination using PCR. 10 μM ROCK inhibitor (Enzo, ALX-270-333-M005) was added after splitting to promote survival. PSC cultures were kept in a humidified atmosphere of 5% $CO_2$ at 37°C and 5% oxygen. All other cultures were kept under atmospheric oxygen condition. Karyotype analysis was performed by MDC Stem Cell Core Facility. Briefly, DNA was isolated using the DNeasy blood and tissue kit (Qiagen, Valencia, CA). SNP karyotyping was assessed using the Infinium OmniExpressExome-8 Kit and the iScan system from Illumina. CNV and SNP visualization were performed using KaryoStudio v1.4 (Illumina).

**Differentiation of NPCs and DA neuronal cultures**

We obtained NPCs and dopaminergic (DA) neurons using a previously published protocol[4]. Briefly, PSCs were detached from Matrigel-coated plates using Accutase (1 mg/ml) and the collected cells were transferred into low-attachment petri dishes and kept for two days in: Neurobasal:DMEM/F12 [1:1], N2 [1x], B27 [1x] with the addition of PMA [0.5 μM], CHIR 99021 [3μM], SB-431542 [10 μM]. From day 2 to day 6, the media was switched to: Neurobasal:DMEM/F12 [1:1], N2 [1x], B27 [1x] with the addition of PMA [0.5 μM], CHIR 99021 [3μM], ascorbic acid [150μM]. On day 6, the suspended cells were transferred onto Matrigel-coated well plates using: Neurobasal:DMEM/F12 [1:1], N2 [1x], B27 [1x]. NPCs were maintained on this media without ROCK inhibitor and used for experiments between passage 7 and 20. For DA differentiation, we used NPCs between passage 7 and 13. To initiate the differentiation, the media was changed to: Neurobasal:DMEM/F12 [1:1], N2 [1x], B27 with vitamin A [1x] with the addition of ascorbic acid [200 μM], FGF8 [100 ng/ml],

PMA [0.5 μM]. After 7 days, the media condition was replaced with: Neurobasal:DMEM/F12 [1:1], N2 [1x], B27 with vitamin A [1x] with the addition of ascorbic acid [100 μM], FGF8 [100 ng/ml], PMA [0.25 μM]. On day 9, cells were split with Accutase and seeded on Matrigel-coated plates in: Neurobasal:DMEM/F12 [1:1], N2 [1x], B27 [1x] with the addition of ascorbic acid [200 μM], cAMP [500 μM], BDNF [10 ng/ml], GDNF [10 ng/ml], and TGFbeta3 [1 ng/ml]. The media was changed every 3-4 days and the differentiated cells were kept in culture for 4 weeks, 6 weeks, and 8 weeks to reach different maturation stages. 10 μM ROCK inhibitor (Enzo, ALX-270-333-M005) was always added after splitting to promote survival.

**CRISPR/Cas9 genome editing**

We prepared CRISPR/eSpCas9 sgRNA plasmid for SVS1C following a published protocol[5]. We first designed a sgRNA sequence (within 10 nucleotides from the target site corresponding to A>G mutation using CRISPOR (http://crispor.tefor.net/). We annealed the oligomer pairs and cloned them into pU6(BbsI)-CAG-Cas9-venus-bpA(#1) plasmid (Addgene ID 86986) carrying eSpCas9 variant from eSpCas9(1.1) plasmid (Addgene ID 71814) with reduced off-target effects and improved on-target cleavage[6]. The PCR product of the eSpCas9 variant were sub-cloned into a CAG expression plasmid (Addgene ID 86986). DNA was submitted for Sanger sequencing to confirm correct sgRNA sequence. We designed 149 nt single-strand oligodeoxynucleotide (ssODN) HDR template to convert G>A (mutation correction) with two silent mutations within sgRNA sequence in the close proximity to the protospacer adjacent motif (PAM) site (3 and 6 nt downstream) to prevent recurrent Cas9 cutting in edited cells. To improve the recombination of ssODN with eCas9-induced double-strand break via single-stranded template repair (SSTR) and ssODN, we applied ectopic expression of RAD52 and of a dominant-negative subfragment of 53BP1 (dn53BP1), which may counteract the endogenous 53BP1[7]. Plasmids encoding components of the DNA repair pathways (human RAD52, and mouse dn53BP1) were kindly obtained from Bruna Paulsen. For the generation of dn53BP1, a fragment containing the tudor domain (residues 1,221 to 1,718 of mouse 53BP1) was sub-cloned and the PCR products of the genes were then sub-cloned into a CAG expression plasmid and sequenced. Transient transfection of plasmids was carried out in SVS1C grown in feeder-free conditions in StemMACS™ iPS-Brew XF culture media (MACS Miltenyi Biotec) in a 6-cell culture plate. One day prior to transfection, we dissociated the cells using Accutase and seeded ~1 x $10^5$ cells per well of a pre-coated 6-well plate as single cells or small clumps. Cells were cultivated in fresh medium containing 10 µM ROCK inhibitor overnight. Lipofection was performed using Lipofectamine 3000 Kit (Thermo Fisher Scientific) according to the manufacturer's protocol. The plasmids were diluted up to 2 mg DNA in 125 ml of Opti-MEM reduced serum medium and added as the DNA-lipid complex to one well of a 6-well plate in a dropwise manner with addition of 5 µM ROCK inhibitor to the culture medium for 24 h. Medium was change on the following day and the cells were kept 48 h in culture until fluorescence-activated cell sorting (FACS). Dissociated cells using Accutase for 5 min were washed and resuspended with DPBS. Then, cells were filtered using Falcon polystyrene test tubes (#352235, Corning) and transferred to Falcon polypropylene test tubes (#352063, Corning). Sorting was performed using BD FACSAria III at the MDC FACS Facility. Sorted cells were suspended in recovery mTeSR™ medium (STEMCELL Technologies) with 1x Penicillin-Streptomycin (P/S) (Gemini Bio-products) and ROCK inhibitor and plated onto 6-well plates (5K cells/well). Growing single cell-derived colonies were transferred from 6-well plates to one well each of 24-well plate and maintained until the colony grew big enough to be partially harvested for DNA isolation using Phire Animal Tissue Direct PCR Kit (Thermo Fisher Scientific) according to manufacturer's protocol. PCR reaction was carried out using 100 ng gDNA in 50 ml with Phusion High-Fidelity DNA Taq polymerase (Thermo Fisher Scientific) according to manufacturer's instructions and annealing temperature of 61˚C. For Sanger sequencing or fragment analysis the PCR products were gel-purified using the Wizard SV Gel and PCR Clean-Up System (Promega). *SURF1* gene product was amplified with SVS1C primers (product length 550 nt). For primer induced restriction analysis (PIRA) the PCR product of 550 nt was cut by *BbsI* (NEB #R3539) into 221+324 nt fragments only in the presence of the mutation c.769G>A. PCR products were submitted to LGC https://www.lgcgroup.com) for Sanger sequencing. Primer sequences, gRNA sequences, and HDR sequence are reported in **Table S5**.

**Generation of iPSC-derived cerebral organoids**

Cerebral organoids were generated according to the protocol described by Lancaster et al., 2014 with some modifications[8]. Shortly, after dissociation into single-cell suspension with Accutase, 10,000 cells were seeded per one well of 96 well-plate in 100 µl of EB medium supplemented with bFGF and 50 µM ROCK inhibitor. After 4 days, medium was replaced with EB medium without bFGF and ROCK inhibitor and at day 6 with neural induction medium (NIM). At day 11, organoids were embedded in Matrigel (Corning, 356234) and kept in NIM for two days, and in organoid differentiation medium without retinoic acid (RA) for another four days. Next, organoids were transferred to ultra-low attachment 6-well plates and culture on orbital shaker (80 rpm) in organoid maturation medium. The composition of the original organoids maturation medium was changed by adding: chemically defined lipid concentrate (1x), ascorbic acid (0.4 mM), BDNF (20ng/ml), HEPES. The organoid size was analyzed by measuring the area using ImageJ software. Each human cerebral organoid was fixed in 4% paraformaldehyde overnight at 4°C, dehydrated by 40% sucrose in PBS and embedded in Tissue-Tek® O.C.T.™ Compound. 12 µm sections were cut and mounted onto slides (Thermo Fisher Scientific). Mounted sections were incubated for 1 h at room temperature with blocking solution (5% normal goat serum + 0.3% Triton X-100 in PBS) and incubated with primary antibodies diluted in blocking solution overnight at 4°C. After three washes with PBST (0.1% TritonX-100), corresponding fluorophore-conjugated secondary antibodies diluted in the blocking solution were added and incubated for 2 h at room temperature and followed by DAPI staining. Finally, stained slides were washed with PBST three times, mounted, and analyzed using a Keyence bz-x710 microscope.

**RNA-sequencing**

PolyA mRNA-seq was carried out for the samples: DA 4w of H1, XM001, SVS1A, SVS1C and DA 8w of H1, XM001, SVS1A, SVS1C (each done in biological triplicates). Total RNA was isolated using the Qiagen RNeasy Mini Kit (#74106, Qiagen) and quality-checked by Nanodrop analysis (Nanodrop Technologies). mRNA-seq was performed by BGI using an oligo dT selection (mRNA enrichment) strategy with oligo dT beads to select mRNA with poly A tail using BGISEQ-500 with DNB seq technology. Ribo-zero total RNA-seq was carried out for the samples: DA 4w of SVS1C and SVS1C.C and DA 8w of SVS1C and SVS1C.C (each done in biological triplicate). 500 ng of total RNA were rRNA was depleted using RNase H-based protocol. Briefly, total RNA was mixed with 1 μg of a DNA oligonucleotide pool comprising 50-nt long oligonucleotide mix covering the reverse complement of the entire length of each mouse rRNA (28S rRNA, 18S rRNA, 16S rRNA, 5.8S rRNA, 5S rRNA, 12S rRNA), incubated with 1U of RNase H (Hybridase Thermostable RNase H, Epicentre), purified using RNA Cleanup XP beads (Agencourt), DNase treated using TURBO DNase rigorous treatment protocol (Thermo Fisher Scientific) and purified again with RNA Cleanup XP beads. rRNA-depleted RNA samples were further fragmented and processed into strand-specific cDNA libraries using TruSeq Stranded Total LT Sample Prep Kit (Illumina) and sequenced on NextSeq 500, High Output Kit, 2 x 76 cycles. Raw sequencing reads were mapped to the human genome (GRCh38 assembly) using STAR (version 2.6.0c) aligner[9]. We used the default settings, with the exception of --outFilterMismatchNoverLmax, which was set to 0.05. Reads were counted using the htseq-count tool, version 0.9.1[10], with gene annotation from GENCODE release 27[11]. Differential gene expression analysis was performed using the DESeq2 (version 1.20.00) R package[12]. For mRNA-seq, read counts for genes expressed in H1, XM001, SVS1A and SVS1C were summed up across the triplicates. SVS1A and SVS1C were treated as disease replicates, and compared to H1 and XM001. For ribo-zero sequencing, SVS1C was compared to the SVS1C.C. All genes with the adjusted P-value lower than 0.05 were considered differentially expressed. Functional enrichment analysis was done using the gProfileR R package[13], version 0.6.6, with default settings. All expressed genes were used as background. All R scripts are available on request. All RNA-seq data have been deposited in the Gene Expression Omnibus (GEO) database (accession number GSE126360).

To review GEO accession GSE126360, enter token enknmcauzbkxdkj into the box:
https://www.ncbi.nlm.nih.gov/geo/query/acc.cgi?acc=GSE126360

**PCR analyses**

Gene expression analysis was performed by quantitative real-time RT-PCR (qPCR) using SYBR Green PCR Master Mix and the ViiA™ 7 Real-Time PCR System (Applied Biosystems). For each target gene, cDNA samples and negative controls were measured in triplicates using 384-Well Optical Reaction Plates (Applied Biosystems). Relative transcript levels of each gene were calculated based on the -ΔΔCT method. Data were normalized to the housekeeping gene *ACTB* and are presented as mean LOG2 ratios in relation to control cell lines. For primer induced restriction analysis (PIRA) of S1A, the PCR product of 141 bp was cut by *SmaI* (NEB #R0141S) into 23 + 118 bp in the presence of c.530T>G mutation. For PIRA of S1C, the PCR product of 437 bp was cut by *AvaII* (NEB #R0153S) into 292+145 bp fragments only in the presence of the mutation c.769G>A. For PIRA of SVS1C, the PCR product of 550 bp was cut by *BbsI* (NEB #R3539) into 221+324 bp fragments only in the presence of the mutation c.769G>A. Primer sequences are reported in **Table S5**.

**Immunostaining**

Cells grown on Matrigel-coated coverslips were fixed with 4% paraformaldehyde (PFA, Science Services) for 20 min at RT and washed two times with PBS. For permeabilization, cells were incubated with blocking solution containing 10% normal donkey serum (DNS) and 1% Triton X-100 (Sigma-Aldrich) in PBS with 0.05% Tween 20 (Sigma-Aldrich) (PBS-T) for 1 h at RT. Primary antibodies were diluted in blocking solution and incubated overnight at 4$^{o}$C on a shaker. Primary antibodies used were as follows: PAX6 (Covance, 1:200), SOX2 (Santa Cruz, 1:100), TUJ-1 (Sigma-Aldrich, 1:3000), OCT4 (Santa Cruz, 1:300), LIN28 (ProteinTech Europe, 1:300), TRA-1-60 (Millipore, 1:200), MAP2 (Synaptic System, 1:100), GFAP (Synaptic Systems, 1:500), NANOG (R&D Systems, 1:200), (SMA) (DakoCytomation, 1:200), SOX17 (R&D Systems, 1:50), TH (Millipore, 1:300), FOXA2 (Sevenhills, 1:100), S100ß (Abcam, 1:500), SYP (Sigma-Alrich, 1:500); VAMP2 (Synaptic Systems, 1:500), NURR1 (Sigma-Aldrich, 1:500). Corresponding secondary antibody (Alexa Fluor, 1:2000, Life Technologies) were diluted in blocking solution for 1 h at RT on a shaker. Counterstaining of nuclei was carried out using 1:10,000 Hoechst (ThermoFisher). All images were acquired using the confocal microscope LSM510 Meta (Zeiss) in combination with the AxioVision V4.6.3.0 software (Zeiss) and further processed with AxioVision software and ImageJ.

**Western blotting**

NPCs were lysed in Lysis Buffer (200 mM NaCl, 50 mM Tris-HCL pH 8.0, 0.05% SDS, 2 mM EDTA, 1% NP40, 0.5% sodium deoxycholate supplemented with (100 mM NaF, 10 µM PMSF, 1:25 EDTA-free protease inhibitor cocktail (Merck), 1:1000 Benzonase (Roche)) for 1 h at 4°C. Protein concentration was determined using the Pierce™ BCA assay (Thermo Fisher Scientific). 120 µg proteins for each samples were loaded on NuPAGE Novex

4-12% Bis-TRIS precast SDS-PAGE gels (ThermoFisher Scientific) and transferred onto Immobilin-FL PVDF membranes (Merck Millipore) by semi-dry transfer. The primary antibodies used were as follows: MT-CO2 (ab110258 Abcam, 1:1,500) and anti-β-tubulin (Sigma, 1:4,000). Chemiluminescence was measured in a Fujifilm LAS-3000 after the addition of Pierce ECL Western Blotting Substrate (Thermo Fisher Scientific).

**COX enzyme activity and histochemistry**

The activities of cytochrome c oxidase (COX) for complex IV and of succinate dehydrogenase (SDH) for complex II were assessed using a colorimetric assay on iPSC-derived NPCs. Enzyme histochemical stains of cytochrome C oxidase and succinate dehydrogenase were performed using standard procedures (Muscle Biopsy, Dubowitz, Sewry, Oldfors, Saunders Elsevier 2013, pp23-24). After gentle centrifugation, NPCs from SVS1C and from genetically corrected SVS1C.C (clones 3p and 3bb) were carefully transferred on Tissue-Tek® on a cork plate and shock frozen in isopentane pre-cooled in liquid nitrogen. 10 μm thick cryosections of these cells were stained by SDH, COX and combined COX-SDH.

**Bioenergetic assessment**

Live-cell assessment of cellular bioenergetics was performed using Seahorse® XF96 extracellular flux analyzer (Seahorse Bioscience), as described previously[2]. Briefly, 20,000 cells were plated into each Matrigel-coated well of the XF96 well plates. NPCs were maintained in the plates for two days, while DA cultures were maintained in the plates for 4 weeks or 8 weeks. On the assay day, the cells were incubated at 37°C 5% $CO_2$ for 60 min to allow media temperature and pH to reach equilibrium before starting the simultaneous measurement of mitochondrial respiration (oxygen consumption rate, OCR) and anaerobic glycolysis (extracellular acidification rate, ECAR) using the sequential introduction of oligomycin, FCCP, and then rotenone plus antimycin A (all products at 1 μM and from

Sigma). Normalization to DNA content in each well of the plate was performed using the CyQUANT Kit (Molecular Probes). The supernatants were stored before and after the seahorse assay and used for lactate measurement using a Lactate Fluorometric Assay Kit (BioVision). Hypoxia and drug treatments were conducted O.N. one day before the assay day. The following treatments were used in this experiment; α-Tocotrienol (AT3) (#10008377, Cayman Chemical), FGF21 (#100-42, PeproTech), glucose, and pyruvate at different concentrations.

**Magnetic-Activated Cell sorting (MACs)**

We performed Magnetic-Activated Cell sorting (MACs) in order to quantify A2B5-positive cells, NCAM-positive cells, GLAST-positive cells, CD11b-positive cells, and O4-positive cells within the 4w and 8w DA neuronal cultures. We used A2B5-APC antibody (130-093-582; Miltenyi Biotec), GLAST (ACSA-1)-APC antibody (130-095-814; Miltenyi Biotec), PSA-NCAM-PE antibody (130–093-274; Miltenyi Biotec), CD11b-VioBlue antibody (130–095-822; Miltenyi Biotec), and O4-APC antibody (130–109-153; Miltenyi Biotec), according to the manufacturer's instructions.

**Cytokine secretion analysis**

The supernatant of the samples used for RNA-Seq (DA 4w of H1, XM001, SVS1A, SVS1C and DA 8w of H1, XM001, SVS1A, SVS1C) were collected and concentrated with Amicon 10K. The samples were assayed in triplicates using the Pro-inflammatory Panel I (MesoScale Discovery, Gaithersburg, MD, USA) for quantitative measurement of 10 cytokines from a single sample volume of 25 μl using an electrochemiluminescent detection method. The cytokines measured were: interferon (IFN)-γ, interleukin (IL)-1β, IL-2, IL-4, IL-6, IL-8, IL-10, IL-12p70, IL-13, and tumor necrosis factor α (TNFα).

**Electrophysiology**

To analyze passive and active membrane properties, spiking and synaptic activity in DA neuronal cultures, whole-cell patch clamp recordings were carried out on week 4, 6 and 8 weeks of differentiation. DA neurons were visualized under phase contrast optics on an upright microscope (Axioskop, Zeiss) by using a 63x/0.95 water immersion objective. Recordings were performed using a patch-clamp amplifier (EPC-9, HEKA Elektronik). Recording pipettes were filled with an intracellular solution containing [mM]: 4 NaCl, 120 KCl, 5 EGTA, 10 HEPES, 5 glucose, 4 $MgCl_2$, 0.5 $CaCl_2$ (pH 7.3, 270 mOsmol/kg). The pipette to bath resistance ranged from 5 to 7 MOhm. Series resistance compensation was applied as much as possible (50–70%). The effective series resistance was in the range of 20–40 MOhm and was checked throughout the whole experiment by using a short depolarizing pulse (10 mV, 20 ms). Recordings were accepted only if the series resistance was less than 40 MOhm. Bath solution contained [mM]: 136 NaCl, 2.5 KCl, 20 glucose, 20 HEPES, 2 $CaCl_2$, 1 $MgCl_2$ (pH 7.3, 305 mOsmol/kg). Whole cell input resistance (RIN) was estimated on the basis of passive current responses to moderate depolarizing voltage pulses of short duration (±10 mV for 20 ms). Whole cell membrane capacitance (CN) was estimated by integration of the capacitive current transient and division by the respective stimulation voltage. Voltage-gated Na+- and K+-currents were elicited by a series of 200 ms depolarizing pulses applied from the holding potential of -70 mV, in 10 mV increments between -70 and +70 mV. Passive responses were subtracted by using a hyperpolarizing pulse of -20 mV. Spontaneous synaptic currents were recorded in voltage-clamp mode at a holding potential of -70 mV without specific blockers. To evaluate action potential generation and discharge properties, cells were adjusted to -90 mV by steady current injection and depolarized by injection of positive current pulses (5-50 pA) of 1 sec duration under current clamp conditions. Signals were acquired at a rate of 10 kHz and analyzed off-line using WinTida 5.0 (HEKA Electronics). All patch clamp experiments were performed at room temperature (20–25°C).

**High-content analysis (HCA)**

HCA-based quantification of neuronal cells and branch complexity was assessed using the CX7 microscope (Thermo Fisher Scientific). Briefly, NPCs or neurons (at 4 weeks or 8 weeks of differentiation) were split using Accutase and seeded at a density of 10,000 cells/well on Matrigel-coated 96 well plates with black-wall and clear-bottom (Corning). The cells were stained with TUJ1 antibody and counter-stained with Hoechst (see below for details on staining method). The morphological changes of TUJ1-postive cells were quantified using the "Cellomics Neuronal Profiling v3.5 BioApplication" (XTI Infinity High Content Platform, Thermo Fisher Scientific). Hypoxia and drug treatments were conducted O.N. one day before the assay day.

**Mitochondrial movement**

DA cultures were grown on 35 mm dishes with coated bottom and 1.5 cover slip. 25 nM MitoTrackerRed CMXRos (Thermo Fischer Scientific) was added for 10 min and then replaced with DA culture media. Live-cell imaging recordings were conducted using a spinning disk microscope (CSU-W, Andor/Nikon) which incubating conditions of 37°C with 5% $CO_2$. We used a 40x oil objective and we imaged the cells every 2 sec. The raw image files were stored as 16-bit in ".nd2" format at 337x337 µm (1024x1024 pixel) at an interval of 2 sec and a total of 200 images per series (total time per series: 6:40 min). The image pre-processing was carried out with Fiji (ImageJ 1.52h) adapted from a previous report[14]. To compensate for photo-bleaching during the time series, we applied the "Bleach Correction" tool, followed by a top-hat spatial filter to increase gray values of mitochondrial objects. The total mitochondria count was calculated by creating a binary image where the minimum gray value (min) was set to min=mean+SD and the maximum gray value (max) was set to max= mean+SD+1, where "mean" and "SD" were obtained by using the "Measure" tool. Next, the “watershed” tool was applied to break up large mitochondria networks. Total number of mitochondria was quantified using the “Analyze Particles” tool with standard setting and a size preference for 8-200 pixel. The moving mitochondria were defined by a particle size of at least 6 pixels that changed location over the time course of 4 frames (=8 seconds). This was accomplished by subtracting the following 4th frame for each frame in the time series (on the top-hat filtered image). Afterwards, the subtracted image series was converted into a binary image series. The number of moving mitochondria was similarly counted using the “Analyze Particles” tool with standard setting and a size preference for 6-200 pixel. Finally, the percentage of moving mitochondria per sample was calculated by dividing the average number of moving mitochondria by the average number of total mitochondria, multiplied with 100. The percentage of stationary mitochondria was obtained by the subtraction of moving mitochondria from 100.

**SURF1 overexpression (OE)**

SURF1 (amino acid sequence NP_003163.1) and mCitrine expressing lentiviral plasmids were generated using the Gateway™ cloning system[15]. Open reading frames from entry vectors encoding the GFP derivative mCitrine or the SURF1 protein (entry clone id: RZPDo839E0486) were shuttled into a lentiviral destination vector harboring a phosphoglycerate kinase (PGK) promoter (pLenti PGK Neo DEST (w531-1) was a gift from Eric Campeau & Paul Kaufman (Addgene plasmid #19067; http://n2t.net/addgene: 19067; RRID: Addgene_19067)[16]. Lentivirus preparation was performed by the Viral Core Facility of the Charité Berlin as previously described[17]. In brief, HEK293T cells were co-transfected with 10 µg of shuttle vector, 5 µg of helper plasmids pCMV-dR8.9, and 5 µg of pCMV-VCV-G using X-tremeGENE 9 DNA transfection reagent (Roche Diagnostics). Cell culture supernatant containing the virus was collected after 72 h and filtered for purification. Virus aliquots were flash-frozen in liquid nitrogen and stored at -80°C. NPCs were seeded on 6 well-plates at a concentration of approximately 500,000 cells per well. The next day, cells were transduced with a mCitrine viruses (GFP) or wild-type SURF1 (SURF1-OE) with a titer of 1.85E+08 and 1.32E+08 particles per ml, respectively. One day later, the medium was changed and NPCs were kept in the incubator for two days to recover. Antibiotic selection was carried out using 500 µg per ml of gentamycin (G418) for 5 days according to killing curves previously performed on NPCs.

**Statistical analysis**

Data were analyzed using GraphPad-Prism software (Prism 4.0, GraphPad Software, Inc.) and R environment for statistical computing. Data presentation and respective statistical analysis of each individual graph are described in the respective figure legends.

#### Supplemental tables and videos

**Table S1. Cytokine profiling in 4w and 8w DA cultures from CTRL ES (H1), CTRL iPS (XM001 and TFBJ), and SURF1 iPS (SVS1A and SVS1C)**

**Table S2. Differentially expressed genes and pathways in 4w DA cultures from SURF1 iPS (SVS1A, SVS1C) vs. WT (H1, XM001) [polyA mRNA-seq]**

**Table S3. Differentially expressed genes and pathways in 8w DA cultures from SURF1 iPS (SVS1A, SVS1C) vs. WT (H1, XM001) [polyA mRNA-seq]**

**Table S4. Differentially expressed genes and pathways in 4w DA cultures and in 8w DA cultures from SVS1C vs. SVS1C.C [ribo-zero total RNAseq]**

**Table S5. List of sequences and primers for qRT-PCR, PIRA, and CRISRP/Cas9 editing**

**Supplemental video 1. Mitochondrial movement in 8w DA cultures from SVS1C.C (pre-processing)**

**Supplemental video 2. Mitochondrial movement in 8w DA cultures from SVS1C.C (post-processing)**

**Supplemental video 3. Mitochondrial movement in 8w DA cultures from SVS1C (pre-processing)**

**Supplemental video 4. Mitochondrial movement in 8w DA cultures from SVS1C (post-processing)**
